# Adenine Base Editing *in vivo* with a Single Adeno-Associated Virus Vector

**DOI:** 10.1101/2021.12.13.472434

**Authors:** Han Zhang, Nathan Bamidele, Pengpeng Liu, Ogooluwa Ojelabi, Xin D. Gao, Tomás Rodriguez, Haoyang Cheng, Jun Xie, Guangping Gao, Scot A. Wolfe, Wen Xue, Erik J. Sontheimer

## Abstract

Base editors (BEs) have opened new avenues for the treatment of genetic diseases. However, advances in delivery approaches are needed to enable disease targeting of a broad range of tissues and cell types. Adeno-associated virus (AAV) vectors remain one of the most promising delivery vehicles for gene therapies. Currently, most BE/guide combinations and their promoters exceed the packaging limit (~5 kb) of AAVs. Dual-AAV delivery strategies often require high viral doses that impose safety concerns. In this study, we engineered an adenine base editor using a compact Cas9 from *Neisseria meningitidis* (Nme2Cas9). Compared to the well-characterized *Streptococcus pyogenes* Cas9-containing ABEs, Nme2-ABE possesses a distinct PAM (N_4_CC) and editing window, exhibits fewer off-target effects, and can efficiently install therapeutically relevant mutations in both human and mouse genomes. Importantly, we show that *in vivo* delivery of Nme2-ABE and its guide RNA by a single-AAV vector can efficiently edit mouse genomic loci and revert the disease mutation and phenotype in an adult mouse model of tyrosinemia. We anticipate that Nme2-ABE, by virtue of its compact size and broad targeting range, will enable a range of therapeutic applications with improved safety and efficacy due in part to packaging in a single-vector system.

## Introduction

Point mutations represent the largest class of known human pathogenic genetic variants [1,2]. Base editors (BEs), which comprise a single-guide RNA (sgRNA) loaded onto a Cas9 (nuclease-inactivated or nickase form) fused to a deaminase enzyme, enable precise installation of A•T to G•C (in the case of adenine base editors (ABEs) [3]) or C•G to T•A (in the case of cytidine base editors (CBEs) [4]) substitutions. In contrast to traditional nuclease-dependent genome editing approaches, base editors do not generate double-stranded DNA breaks (DSBs), do not require a DNA donor template, and are more efficient in editing non-dividing cells, making them attractive agents for *in vivo* therapeutic genome editing.

While robust editing has been achieved in many cultured mammalian cell systems, safe and effective *in vivo* delivery of base editors remains a major challenge. To date, both non-viral and viral delivery methods have shown great promise for delivering base editors for *in vivo* therapeutic purposes in rodents and primates [5,6]. For example, *in vivo* delivery via AAVs has achieved efficient editing in a wide range of tissue and cell types including liver [5–10], heart [11], muscle [12,13], retina [14,15], inner ear [16], and central nervous system (CNS) [17,18]. However, the large coding size (5.2 kb) of the best-characterized *Streptococcus pyogenes* Cas9 (SpyCas9)-containing BEs exceed the packaging limit of AAV (5 kb) [19,20]. Currently, *in vivo* delivery of base editors by AAV has been approached by splitting the SpyCas9 base editor between two AAVs and relying on the use of intein trans-splicing for the assembly of the full-length effector [13,16–18,21,22]. Although effective, this approach requires transduction of the target cell by both AAVs and successful *in trans* splicing of the two intein halves. The requirement to deliver two AAV vectors also increases the viral dosage needed for a treatment, which raises safety concerns and adds burdens to AAV manufacturing [23–25].

Compact Cas9 orthologs are ideal candidates for engineering base editors suitable for single-AAV delivery [26–28]. For example, single-AAV delivery of a domain-inlaid *Staphylococcus aureus* Cas9 (SauCas9) ABE has been reported in cultured HEK293 cells [29]. Previously, we developed a *Neisseria meningitidis* Cas9 (Nme2Cas9) as an *in vivo* genome editing platform [27,30]. Nme2Cas9 is a compact, naturally accurate genome editor with a distinct N_4_CC PAM specificity. Recently, two other groups have successfully implemented Nme2-CBEs in cultured mammalian cells and in rabbits [31], as well as Nme2-ABEs in rice [32]. Here, we develop ABEs using Nme2Cas9 (Nme2-ABEs) and define their editing efficiencies, editing windows, and off-target activities in comparison to those of the widely applied Spy-ABEs in cultured mammalian cells. Next, we show that Nme2-ABE can edit multiple therapeutically significant loci, including one of the most common mutations occurring in Rett syndrome patients that cannot be targeted by other compact ABEs (e.g. ABEs derived from SauCas9 and SauCas9-KKH [33–36]) due to PAM restrictions. Lastly, by optimizing the promoter and the nuclear localization signals, we show that Nme2-ABE and its guide can be packaged into a single-AAV vector genome for *in vivo* delivery. One systematic administration of the single-AAV vector encoding both Nme2-ABE and sgRNA readily corrects the disease-causing mutation and phenotype in an adult mouse model of hereditary tyrosinemia type 1 (HT1).

## Results

### Development of Nme2-ABE and comparison of editing windows and off-target effects to those of Spy-ABE

First, to evaluate base editing efficiency in a streamlined manner, we developed an ABE reporter cell line in which a G-to-A mutation in the mCherry coding sequence generates a nonsense mutation. Adenine base editing can reverse the mutation and recover red fluorescence, and the editing efficiency can be readily measured by fluorescence-activated cell sorting (FACS) (**Figure 1a, Supplementary Figure 1**). Initially, we constructed Nme2-ABE7.10 by linking a TadA-TadA7.10 dimer from the Spy-ABE7.10 to the N-terminus of the Nme2Cas9 HNH nickase [3]. However, by plasmid transient transfection, Nme2-ABE7.10 showed poor activity in the ABE reporter cell line. Because the evolved TadA8e is highly active and compatible with a wide range of Cas9s [35], we next engineered Nme2-ABE8e by linking TadA8e monomer to the N-terminus of the Nme2Cas9 HNH nickase (**Figure 1b**). We found that Nme2-ABE8e supports robust editing activity in the ABE reporter cell line (**Figure 1c**). Next, to define the editing window and editing efficiency of Nme2-ABE8e, and to compare them to those parameters of Spy-ABE7.10 and Spy-ABE8e, we transfected HEK293T cells with plasmids expressing each ABE along with sgRNAs targeting 12 human genomic loci for Nme2-ABE8e [including eight dual-target sites (target sites followed by NGGNCC PAMs for both SpyCas9 and Nme2Cas9) [30] and four Nme2Cas9-specific target sites]. We found that this first-generation Nme2-ABE8e has a broad but shallow editing window that, due to modest activity, was difficult to define clearly (**Figure 1d**).

**Figure 1.**
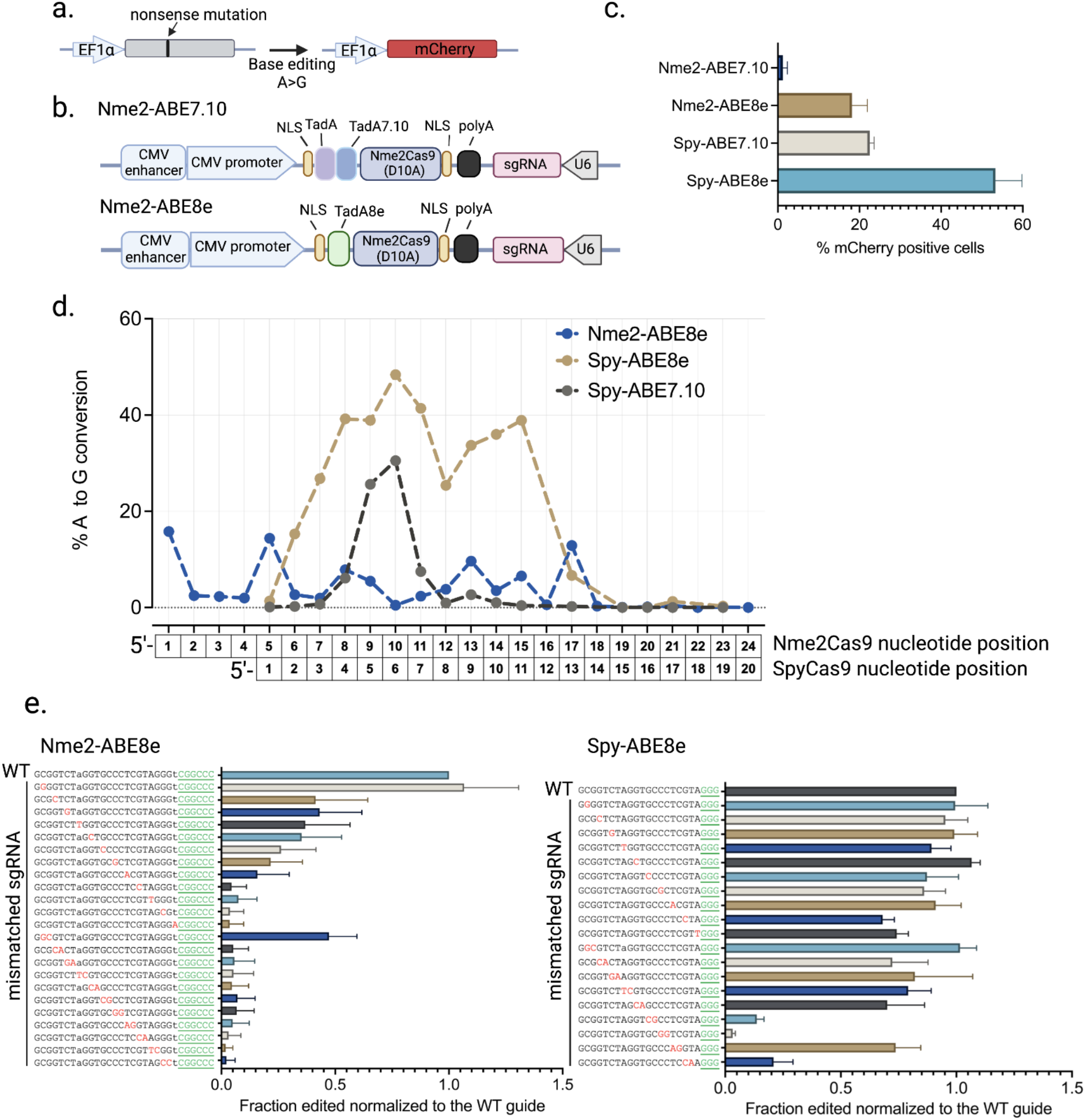
**a).** Schematic representation of the ABE reporter HEK293T cell line. **b).** Schematic representation of the Nme2-ABE constructs. **c).** Comparison of editing efficiencies of Nme2-ABEs to those of Spy-ABEs in the ABE reporter cell line by plasmid transient transfection (n = 3 biological replicates). **d).** Summary of editing windows and comparison of editing efficiencies for the ABEs at endogenous genomic loci. Each data point represents the mean A-to-G editing efficiency at the indicated position of the spacer across 12 Nme2Cas9 target sites and 8 SpyCas9 target sites, respectively (n = 3 biological replicates). Summary of the individual A-to-G conversion and indel efficiencies at each target site can be found in **Supplementary Figure 2**. **e).** Comparison of Nme2-ABE8e mismatch tolerance to that of Spy-ABE8e in the ABE reporter cell line. The activities of the effectors with the mismatched guides are normalized to that of the perfectly complementary (WT) guide. Red, mismatched nucleotides; green, PAM sequence (n = 3 biological replicates).

We next sought to understand the potential extent of off-target editing by Nme2-ABE8e. It has been shown that the major source of DNA off-target base editing is Cas9-dependent [37,38], caused by Cas9 binding and unwinding at near-cognate sequences. Because Nme2Cas9 is highly accurate during nuclease-driven editing in cells and *in vivo* [27,30,39], we hypothesized that Nme2-ABE8e would exhibit similar accuracy advantages relative to Spy-ABE8e. As an initial test, we systematically investigated the tolerance for nucleotide mismatches between the guide and the target sequence for the two effectors. We designed a panel of guides targeting the ABE reporter with single- and di-nucleotide mismatches with the target sequence for both Nme2-ABE8e and Spy-ABE8e and measured their activities by plasmid transfection and FACS (**Figure 1e**). Considering the differences in on-target efficiencies between the two effectors, we further normalized the activities of the mismatched guides to those of the perfectly complementary guides for each effector. We found that Nme2-ABE8e exhibited significantly lower off-target editing propensity than Spy-ABE8e: while single-nucleotide mismatches in the seed region (guide nucleotide positions 17-24 for Nme2Cas9, and 10-20 for SpCas9) and the majority of dinucleotide mismatches significantly compromised the editing efficiency of Nme2-ABE8e, these near-cognate sequences were mostly efficiently edited by Spy-ABE8e (**Figure 1e**).

### Inhibition of base editing by anti-CRISPR proteins that limit DNA binding activity

The development of Nme2Cas9 base editing platforms raises the possibility that regulation strategies developed for nuclease-based editing could be similarly implemented in the case of base editing. One such strategy is the use of anti-CRISPR (Acr) proteins that limit Cas9 DNA-binding activity [40–43], which have been deployed to reduce both off-target [44] and off-tissue editing [45]. AcrIIC3 and AcrIIC4 have been reported to reduce Nme2Cas9 DNA binding activity [41,43,46] but have no effect on SpyCas9 [41,43]. To test whether such Acr proteins can function as off-switches for Nme2-ABE8e base editing, we co-transfected the ABE reporter cell line described above with plasmids expressing Nme2-ABE8e, sgRNA, and Acr proteins. Spy-ABE7.10 and AcrIIA4 (an anti-CRISPR that prevents SpyCas9 DNA binding [40], nuclease editing [40,44], and base editing [47]) was used as a positive control. Conversely, AcrE2, which is a Type I-E Acr [43] that has no effect on SpyCas9 or Nme2Cas9 activity, was used as a negative control. As expected, Spy-ABE7.10 base editing was reduced to background levels by AcrIIA4, but AcrE2, AcrIIC3, and AcrIIC4 had no effect (**Supplementary Figure 3a**). By contrast, Nme2-ABE8e editing was strongly inhibited by AcrIIC3 and AcrIIC4, but AcrE2 and AcrIIA4 had no effect. These results confirm that these anti-CRISPRs can be effective off-switches for Nme2-ABE8e editing.

Tissue-specific miRNAs, in combination with miRNA response elements [MREs] in the 3’UTR of an Acr construct, have been used to restrict editing to cell types that express such miRNAs, both in cultured cells [41,48] and *in vivo* [45]. To determine whether such a strategy could be used to control Nme2-ABE8e, we inserted miR-122 MREs into the 3’UTRs of our AcrIIC3 and AcrIIC4 constructs. Mir-122 is a hepatocyte-specific miRNA that is expressed in Huh7 cells but not HEK293 cells, and has been used to validate this strategy for nuclease-based editing. Again, AcrIIC3 and AcrIIC4 inhibited Nme2-ABE8e activity at an endogenous site in HEK293T cells, and inhibition largely persisted even with the MREs (**Supplementary Figure 3b**), as expected since miR-122 is not present to silence Acr expression. In Huh7 cells, which are transfected with lower efficiencies, AcrIIC3 and AcrIIC4 again inhibited Nme2-ABE8e, but this inhibition of editing activity was largely relieved by the insertion of the MRE122 sites. These results indicate that miRNA-repressible anti-CRISPRs can be used to enforce the cell-type specificity of base editing, as it can for nuclease editing.

### Installation of therapeutically relevant edits with Nme2-ABE8e

We next tested the potential of Nme2-ABE8e to correct pathogenic mutations or to introduce disease-suppressing mutations. One of the most common mutations that cause Rett syndrome is a C•G to T•A base transition that produces a nonsense mutation in the human *MeCP2* gene (c.502 C>T; p.R168X) [49–51]. Because the target adenine is within a cytidine-rich region, this mutation is not readily accessible for ABEs based on other compact Cas9s such as SauCas9 and SauCas9-KKH [33–36]. To test whether Nme2-ABE8e can correct this pathogenic mutation, we electroporated Nme2-ABE8e mRNA with a synthetic sgRNA into a Rett syndrome patient-derived fibroblast cell line that possesses this mutation. By amplicon deep sequencing, we found that Nme2-ABE8e successfully edited the target adenine (A10) (**Figure 2a, 2b**). A bystander edit at an upstream adenine (A16) causes a missense mutation (c.496 T>C; p.S166P), although this occurs with only one-fourth the frequency of intended edit at A10. Because S166 has been shown to be subject to phosphorylation in mice and is conserved from *X. laevis* to humans [52], further investigation will be needed to determine whether bystander editing at A16 impairs functional rescue of edited cells. However, as with other base editors [3,29,53–58], future protein engineering efforts adjusting the editing window promise to provide greater control over Nme2-ABE editing outcomes.

**Figure 2.**
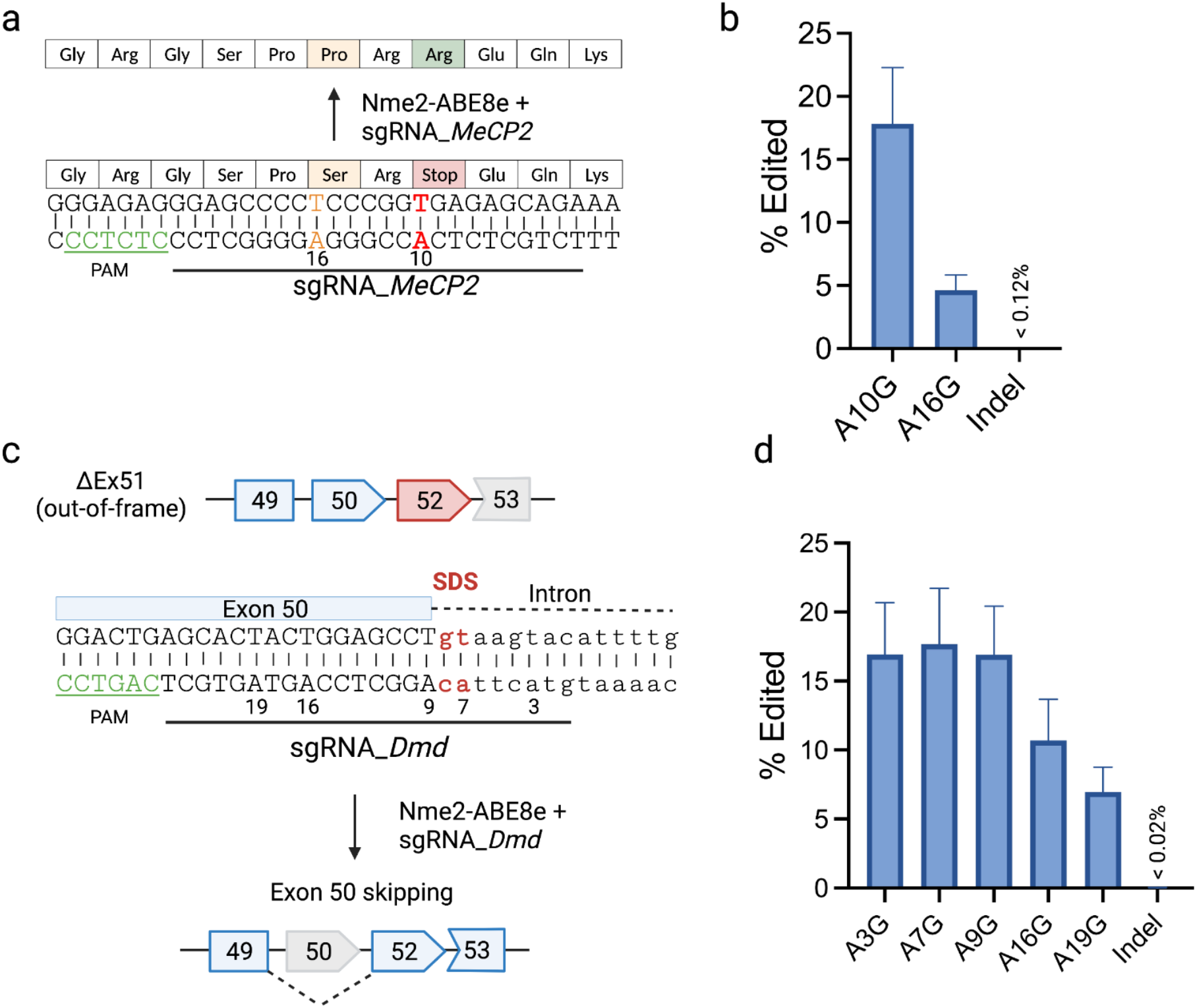
**a).** Schematic representation of a nonsense mutation in the human *MeCP2* gene (c.502 C>T; p.Rl68X) that causes Rett Syndrome. The black underline denotes the target sequence of an Nme2-ABE8e guide for reverting the mutant A to G (wildtype) at position 10 (red, bold). The PAM region is underlined in green. A bystander edit at position 16 (orange) can generate a missense mutation (c.496 T>C; p.S166P). **b).** Amplicon deep sequencing quantifying the editing efficiency in Rett patient-derived fibroblasts transfected with the Nme2-ABE8e mRNA and the synthetic sgRNA_*MeCP2* noted in **a** (n = 3 biological replicates). **c).** Schematic representation of the exon skipping strategy that restores the reading frame of the mouse *Dmd* transcript. Deletion of exon 51 (ΔEx51) can alter the reading frame and generate a premature stop codon in exon 52 (red). Adenine base editing at the splice site of exon 50 (red) by Nme2-ABE8e can cause exon 50 skipping (gray) and restore the *Dmd* reading frame. The PAM region is underlined in green. **d).** Amplicon deep sequencing quantifies the editing efficiency at the target site in mouse N2a cells transfected with the Nme2-ABE8e and sgRNA_*Dmd* expression plasmid (n = 3 biological replicates).

Next, we sought to generate a disease-suppressing mutation that has been shown to reverse phenotypes of a validated Duchenne muscular dystrophy (DMD) mouse model (ΔEx51) [59]. The ΔEx51 mouse model was generated by deletion of the exon 51 in the *Dmd* gene, resulting in a downstream premature stop codon in exon 52, causing the production of a nonfunctional truncated dystrophin protein. Previously, it has been shown that the *Dmd* reading frame can be restored by skipping exon 50 by adenine base editing (**Figure 2c**) [13]. However, *in vivo* base editing using ABEmax-SpCas9-NG delivered by dual-AAV vectors was limited to local muscle injection due to the high viral dosage required to achieve therapeutic benefit. We identified a guide design for Nme2-ABE8e to target the adenine (A7) within the splice donor site downstream of exon 50 (**Figure 2c**). By plasmid transfection in the mouse N2a cell line and amplicon deep sequencing, we found that Nme2-ABE8e can generate 17.67 ± 4.57% editing at A7 (**Figure 2d**). The efficient editing at multiple bystander adenines is not a concern in this case as those adenines are within the skipped exon 50 or the intron.

### Optimization of an Nme2-ABE8e construct for single AAV delivery

Previously, we showed that Nme2Cas9 with one or two sgRNAs can be packaged into a single AAV vector and support efficient editing *in vivo* [27,30]. We reasoned, based upon the compact sizes of Nme2Cas9 and TadA8e, that Nme2-ABE8e with a sgRNA could be packaged into a single AAV for *in vivo* delivery. To achieve this, we first replaced NmeCas9 with Nme2-ABE8e in the minimized all-in-one AAV vector reported previously [27]. We attached one cMyc NLS sequence on each terminus of Nme2-ABE8e while retaining the original promoters for effector and sgRNA expression (**Figure 3a,** 2x cMyc). By transient transfection of vector backbone plasmids, the single-AAV construct successfully edited the ABE reporter cell line. To further improve Nme2-ABE editing efficiency, we then tested three different NLS configurations: 1) one cMyc NLS on the N-terminus and two cMyc NLSs on the C-terminus (3x cMyc); 2) one Ty1 NLS, which derived from the yeast Ty1 retrotransposon that supports robust nuclear localization in dPspCas13b fusion proteins, on the N-terminus (Ty1) [60]; and 3) one bipartite SV40 NLS (BP_SV40) on each terminus (2x BP_SV40) (**Figure 3a**) [61]. When transfecting the vector plasmid into the ABE reporter cell line, the 2x BP_SV40 construct showed the highest editing efficiency (**Figure 3b**).

**Figure 3.**
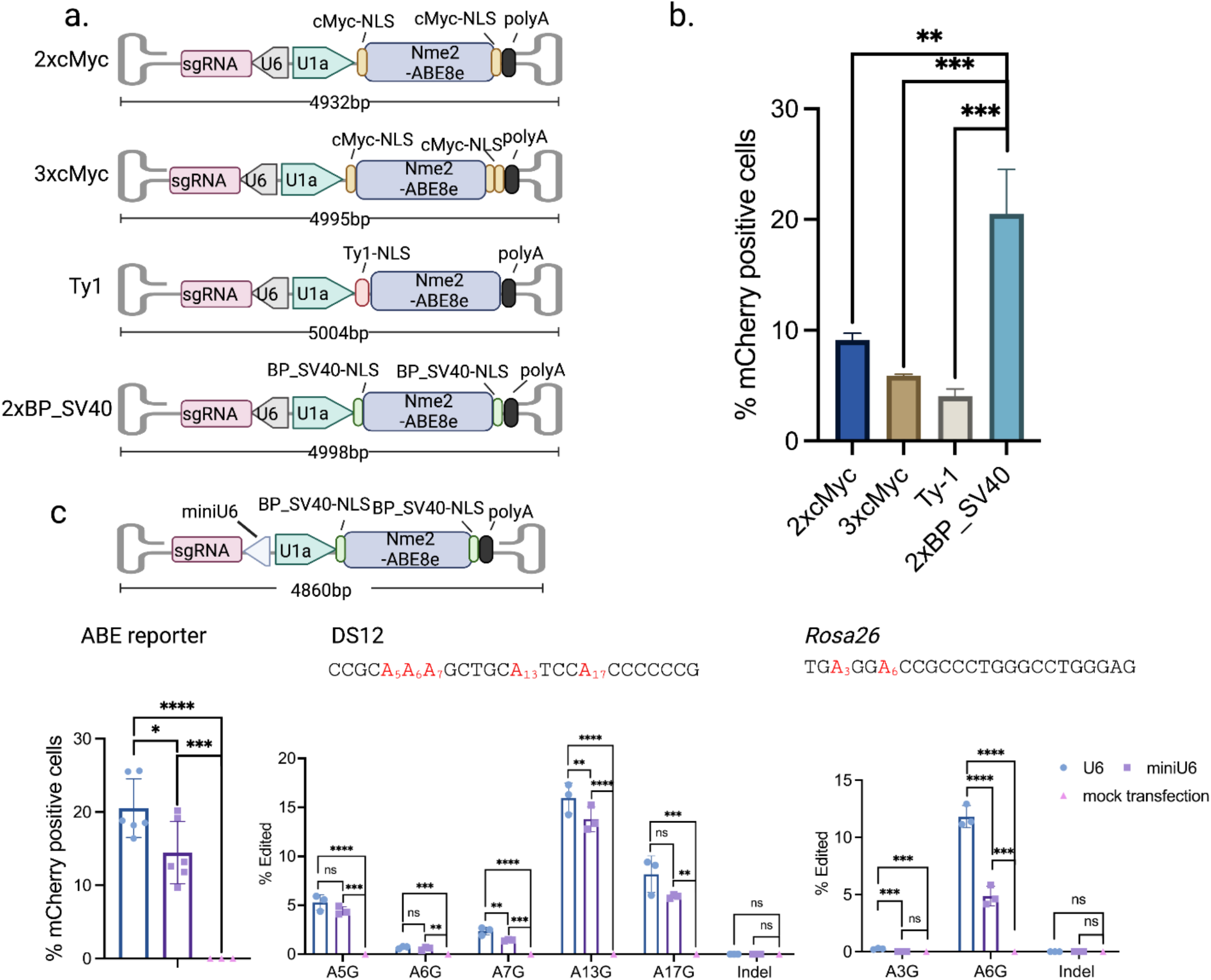
**a).** Schematic representation of single AAV constructs with different NLS configurations. **b).** Comparison of different NLS configurations by plasmid transfection in the ABE reporter cell line. **c).** Comparison of Nme2-ABE activity when sgRNA expression is driven by the U6 or miniU6 promoters in the 2xBP_SV40 NLS construct targeting the ABE reporter site (left) or endogenous human (middle) and mouse (right) genomic sites by plasmid transfection in cultured cells followed by amplicon deep sequencing (n = 3 biological replicates). Data represent mean ± SD; ns, P > 0.05; *, P < 0.05; **, P < 0.01; ***, P < 0.001; ****, P < 0.0001 (one-way ANOVA).

The total length of the vector constructed with the 2xBP_SV40 NLS, hereafter Nme2-ABE8e-U6, is 4998 bp, very close to the packaging limit of AAV. To test whether we could further reduce the vector size without significantly compromising editing efficiency, we turned to a recently reported “miniU6” promoter that has been shown to support sgRNA expression and achieve comparable editing efficiencies as with the complete U6 promoter [62]. Upon replacement of the U6 promoter with miniU6 promoter, the vector, hereafter Nme2-ABE8e-miniU6, was shortened to 4860 bp, within the packaging limit of AAV (**Figure 3c**). Both constructs induced robust editing in the ABE reporter cell line via transient transfection of the vector plasmids (**Figure 3c**).

To avoid potential ABE reporter-specific effects, we further tested both single-AAV vector backbone plasmids at two endogenous target sites: 1) one of the human dual-target sites, DS12, and 2) a previously reported Nme2Cas9 target site in the mouse *Rosa26* gene [45]. By plasmid transfection in human HEK293T or mouse N2a cells, we observed significant editing at these loci by both vectors, although the Nme2-ABE8e-miniU6 vector was somewhat less efficient (**Figure 3c**). We thus chose both vector designs for the subsequent *in vivo* study.

### Hydrodynamic injection of single-AAV vector plasmids corrects the disease mutation and phenotype in an adult mouse model of HT1

To test the *in vivo* editing efficiency and therapeutic potential of the single-AAV constructs, we chose to target a pathogenic mutation associated with the liver disease HT1. HT1 is caused by mutations in fumarylacetoacetate hydrolase (FAH), which catalyzes one step of the tyrosine catabolic pathway. FAH deficiency leads to accumulations of toxic fumarylacetoacetate and succinyl acetoacetate, causing liver, kidney, and CNS damage [63]. The *Fah*^PM/PM^ mouse model possesses a G•C to A•T point mutation in the last nucleotide of exon 8, which causes skipping of exon 8 and FAH deficiency (**Figure 4a**). Without treatment, FAH deficient mice will rapidly lose weight and eventually die. The *Fah*^PM/PM^ mouse can be treated with 2-(2-nitro-4-trifluoromethylbenzoyl)-1,3-cyclohexanedione (NTBC), an inhibitor of an enzyme upstream within the tyrosine degradation pathway, which prevents toxin accumulation [64]. Previously, we and others have tested various *in vivo* gene-editing tools to treat the *Fah*^PM/PM^ mouse model, including Cas9-directed HDR [27,65,66], base editing [67,68], microhomology-directed end joining [69], and prime editing [70]. Among these, multiple approaches including AAV, lipid nanoparticle (LNP), and plasmid hydrodynamic tail-vein injection have been used to deliver the gene-editing agents into this mouse model. However, caveats should be considered when comparing the efficiencies of different gene-editing strategies: only the initial editing efficiency (measured before NTBC withdrawal) reflects the activity of the gene-editing agents, because after NTBC withdraw, the hepatocytes in which *Fah* has been repaired will show clonal expansion over time due to their survival advantage.

**Figure 4.**
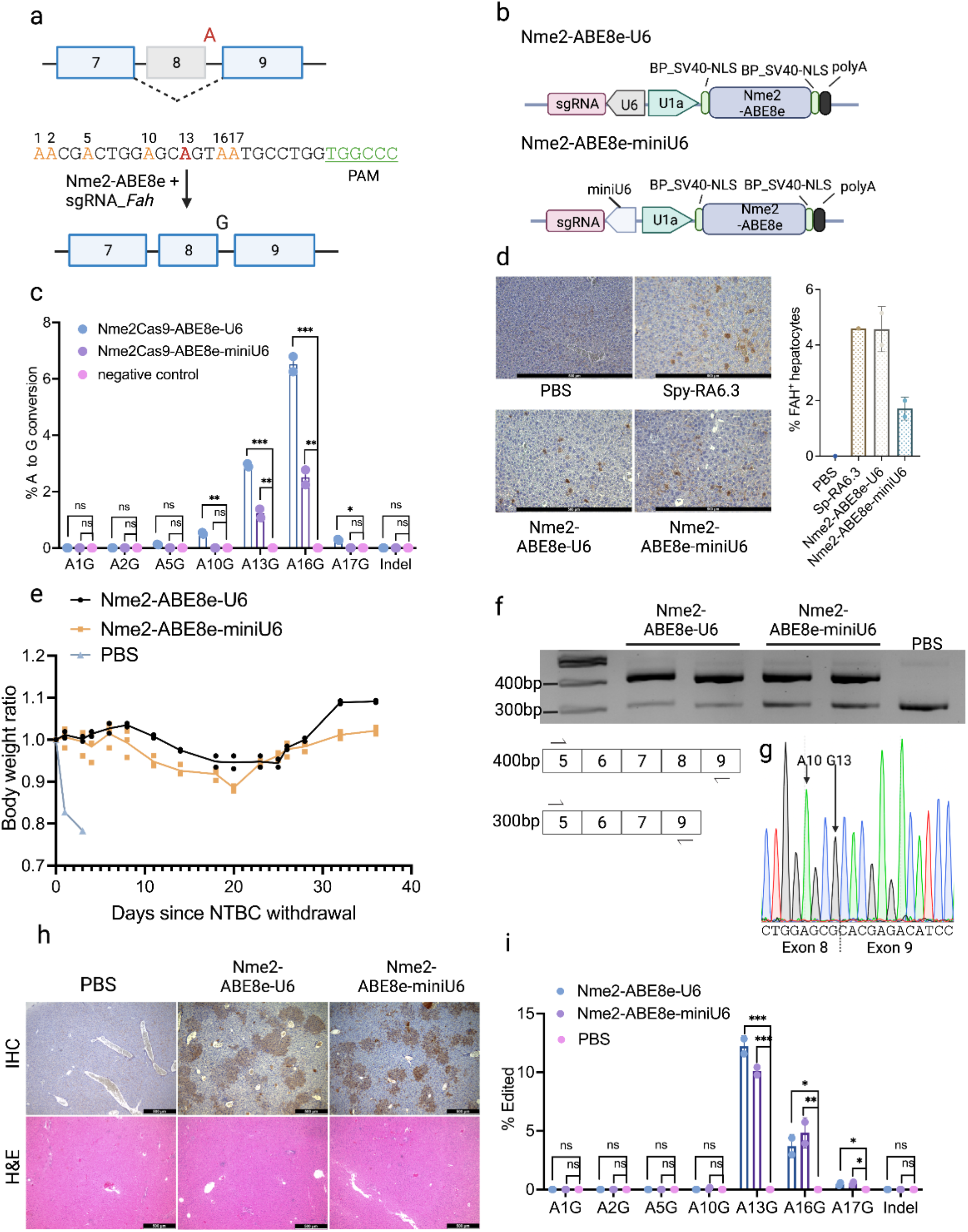
**a).** Illustration of the pathogenic point mutation in the *Fah*^PM/PM^ mouse model that causes exon 8 skipping of the *Fah* gene, and the guide design for Nme2-ABE8e to correct the point mutation. Red and bold, target adenine; orange, other bystander adenines; green and underlined, PAM. **b).** Illustration of constructs of the single-AAV vector plasmids used in *in vivo* studies. **c).** Editing efficiencies at the *Fah* mutant site by AAV plasmid electroporation in MEF cells derived from the *Fah*^PM/PM^ mouse. Data are from amplicon deep sequencing (n = 2 biological replicates). **d).** Anti-FAH immunohistochemistry (IHC) staining showing FAH^+^ hepatocytes, before NTBC withdrawal, in the *Fah*^PM/PM^ mouse hydrodynamically injected with the indicated plasmid. The bar graph quantifies the percentage of FAH^+^ hepatocytes detected by IHC (n = 2 mice per group). Scale bars, 500 μm. **e).** Body weight plot of mice injected with the single-AAV vector plasmid showing gradual weight gain over a month after NTBC withdrawal. **f).** RT–PCR analysis of the plasmid- or PBS-injected mouse livers using primers that span exons 5 and 9. The wild-type amplicon is 405 bp and exon 8 skipped amplicon is 305 bp. **g).** Representative Sanger sequencing trace of the 405 bp RT-PCR band. **h).** Anti-FAH IHC staining showing expansion of FAH^+^ hepatocytes 40 days post NTBC withdrawal. Scale bars, 500 μm. **i).** Quantification of the editing efficiency by amplicon deep sequencing of genomic DNA of the treated mouse livers harvested 40 days post NTBC withdrawal. **d-i**, n = 2 mice per group). Data represent mean ± SD; ns, P > 0.05; *, P < 0.05; **, P < 0.01; ***, P < 0.001 (one-way ANOVA).

We first validated a guide targeting the point mutation by electroporation of the single-AAV vector plasmids, either Nme2-ABE8e-U6 or Nme2-ABE8e-miniU6, into mouse embryonic fibroblasts (MEFs) isolated from a *Fah*^PM/PM^ mouse (**Figure 4b**). We detected modest but significant editing by both vectors at the target adenine at position 13 (A13), despite low (~12%) plasmid electroporation efficiencies. We also observed higher levels of bystander editing at A16 with both vectors, and a lower level of bystander editing at A10 for Nme2-ABE8e-U6. Bystander editing at A10 (which changes an active-site-proximal serine into an alanine) has been observed previously with Spy-ABE as well [67]. The effect (if any) of the intronic A16 edit on intron 8 splice donor activity has not been defined (**Figure 4c**).

To test our single-AAV vectors of Nme2-ABE8e *in vivo*, we first performed hydrodynamic tail-vein injections of the AAV-vector plasmids into 10-week-old HT1 mice, or PBS injection as a negative control group [71]. We also injected plasmids expressing Spy-RA6.3, which is a codon-optimized Spy-ABE, as a positive control [67]. Seven days post-injection, we sacrificed 2 mice from each experimental group, and 1 mouse from each control group, to measure the editing efficiency before hepatocyte expansion. We then withdrew NTBC for the rest of the mice for long-term phenotypic study. Before NTBC withdrawal, anti-FAH immunohistochemistry (IHC) staining showed 4.58 ± 1.1% FAH^+^ hepatocytes from the group that was injected with the Nme2-ABE8e-U6 plasmid, and 1.71 ± 0.49% from the group injected with the Nme2-ABE8e-miniU6 plasmid (**Figure 4d**). The mouse injected with Spy-RA6.3 plasmid showed 4.5% FAH^+^ hepatocytes, consistent with previously reported data [67] (**Figure 4d**). After NTBC withdrawal, we monitored body weight changes. The PBS injected mice rapidly lost body weight after NTBC withdrawal and were euthanized. By contrast, mice injected with either the Nme2-ABE8e-U6 or Nme2-ABE8e-miniU6 plasmid gradually gained body weight, suggesting rescue of the pathological phenotype (**Figure 4e)**. Forty days after NTBC withdrawal, we sacrificed mice from all surviving groups. To determine whether Nme2-ABE8e successfully corrects the *Fah* splicing defect, we extracted total RNA from the livers and performed reverse transcription PCR (RT-PCR) using primers that spanned exons 5 and 9. By contrast to the PBS-injected mice, which only showed a 305 bp PCR product corresponding to the truncated mRNA lacking exon 8, we observed that the 405 bp PCR product (containing exon 8) predominated in the Nme2-ABE treated mice **(Figure 4f)**. Sanger sequencing of the 405 bp bands further confirmed the presence of the corrected G residue at the end of exon 8 **(Figure 4g)**. When performing anti-FAH IHC staining, we observed expansion of FAH^+^ hepatocytes in the groups that were injected with either of the single-AAV vector plasmids **(Figure 4h)**. Amplicon deep sequencing of genomic DNA from the livers of treated mice again provided evidence for Nme2-ABE activity **(Figure 4i)**. By contrast to the efficiencies achieved in the MEF cells, we observed a lower editing at A16 and no significant editing at A10, likely due to partial (A16) or complete (A10) selection against mice harboring bystander edits at those positions. These data indicate that our Nme2-ABE8e single-AAV vector plasmids can correct the disease genotype and phenotype of the *Fah*^PM/PM^ mice *in vivo*.

### *In vivo* base editing by single AAV delivered Nme2-ABE8e in the *Fah*^PM/PM^ mice

Encouraged by the initial results, we packaged AAV9 with the Nme2-ABE8e-U6 construct, as well as the Nme2-ABE8e-miniU6 construct, considering that the relatively smaller size of the latter may potentially benefit packaging efficiency **(Figure 5a)**. However, both constructs yielded similar vector titers. Next, to confirm the AAV genome integrity, we performed AAV genomic DNA extraction and alkaline gel electrophoresis. We did not observe any sign of genome truncation **(Supplemental Figure 4)**. We then tail-vein injected 8-week-old *Fah*^PM/PM^ mice at a dosage of 4 x 10^11^ vg per mouse. We kept the mice on NTBC for one month before analyzing the editing efficiency. One month after AAV injection and before NTBC withdrawal, we sacrificed the mice and performed IHC staining using an anti-FAH antibody. The negative control groups injected with AAV9 expressing Nme2-ABE and a sgRNA targeting the *Rosa26* gene did not show any FAH^+^ hepatocytes. In contrast, we observed 6.49 ± 2.08% FAH^+^ hepatocytes in the AAV9-Nme2-ABE8e-U6-*Fah* treated group, and 1.62 ± 0.49% FAH^+^ hepatocytes in AAV9-Nme2-ABE8e-miniU6-*Fah* treated group **(Figure 5b)**. Because repair of 1/100,000 hepatocytes was reported to rescue the phenotype, both AAV constructs achieved efficiencies that were well above the therapeutic threshold [72]. Moreover, the percentage of edited hepatocytes by AAV9-Nme2-ABE8e-U6-*Fah* was higher than what has been reported previously by other genome editing strategies [65–67,70]. By targeted deep sequencing, the editing efficiency at the target adenine (A13) in the AAV9-Nme2-ABE8e-U6 treated group is 0.34 ± 0.14%, while no significant editing was observed in the AAV9-Nme2-ABE8e-miniU6 treated group, possibly due to low efficiency that was below the detection limit of amplicon deep sequencing **(Figure 5c)**. The reason for the higher frequency of FAH^+^ hepatocytes than the frequency of editing at the DNA level is likely due to hepatocyte polyploidy [73], as well as the presence of genomic DNA from nonparenchymal cells. Similar distinctions in FAH^+^ frequencies and genomic readouts were also observed in previous studies using this mouse model [67,70]. We also measured the editing efficiency at the *Rosa26* locus and observed ~5% editing efficiency at the target site, indicating that the efficiency of AAV9-delivered Nme2-ABE8e is target site dependent, and higher-efficiency sites can be identified **(Figure 5d)**. We did not detect any above-background level of indel, indicating that single-AAV delivered Nme2-ABE8e can install precise editing *in vivo* without generating unwanted indels (**Figure 5c, d**).

**Figure 5.**
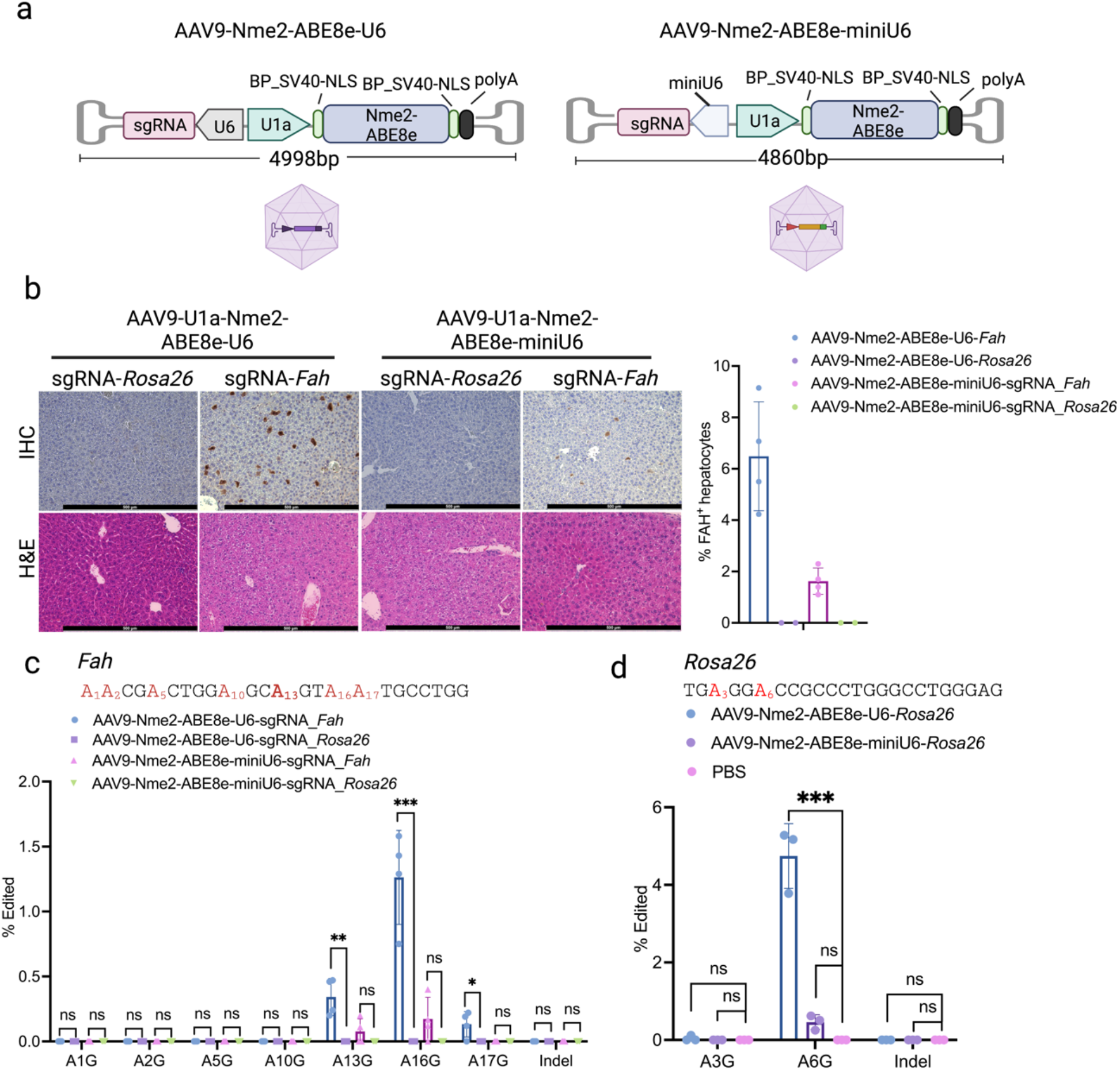
**a).** Schematic representation of the AAV constructs. **b).** Anti-FAH immunohistochemistry (IHC) staining showing FAH^+^ hepatocytes, before NTBC withdrawal, in the *Fah*^PM/PM^ mouse injected with AAV9 expressing Nme2-ABE8e with a sgRNA targeting either the *Fah* gene, or the *Rosa26* gene that serves as a negative control. Scale bar, 500 μm. The bar graph quantifies the percentage of FAH^+^ hepatocytes detected by IHC (n = 4 mice in the *Fah* targeting group, n = 3 mice in the *Rosa26* targeting group). **c, d).** Quantification of the editing efficiency by amplicon deep sequencing of genomic DNA from the AAV9-injected mouse livers harvested before NTBC withdrawal (n = 4 mice in the *Fah* targeting group, n = 3 mice in the *Rosa26* targeting group and the PBS control group). Data represent mean ± SD; ns, P > 0.05; *, P < 0.05; **, P < 0.01; ***, P < 0.001 (oneway ANOVA).

Nme2Cas9 is known to be highly accurate in nuclease based editing in cells [27,30,39], and our mismatch scanning experiments (**Figure 1e**) strongly suggest that the same will be true *in vivo*. To evaluate potential Cas9-dependent off-target effects in the AAV9-injected mice, we searched for genome-wide off-target sites for Nme2Cas9 using Cas-OFFinder [74], allowing for up to 6 mismatches. We then performed amplicon deep sequencing in the AAV9 treated livers at the two top-ranking potential off-target sites, each including 5 mismatches. We did not detect any above-background A•T-to-G•C editing at these sites **(Supplementary Figure 5)**.

### Optimizing effector and sgRNA arrangement improves editing efficiency by AAV delivery

Previous studies have shown that effector and sgRNA placement and orientation within the AAV genome can affect transgene expression levels and editing efficiencies [18, 26, 85]. To test if different arrangements of sgRNA and effector cassettes in the Nme2-ABE8e all-in-one AAV construct can further increase *in vivo* editing efficiency, we moved the U6-sgRNA cassette to the 3’ end of the AAV genome and reversed its orientation, similar to the optimal arrangement reported by Fry *et al*. [85] (**Figure 6a**). Using the same *Rosa26* guide described above (**Figure 5d**), we packaged the rearranged construct in AAV9 capsids and performed tail-vein injections in 8-week-old mice. We observed significantly improved editing efficiency (34 ± 11.6%) with the optimized construct (**Figure 6b**), which is significantly greater than the 4.7 ± 0.94% efficiency with the original configuration, and comparable to the editing efficiency achieved previously by dual-AAV delivered split-intein SpyCas9-ABEmax targeting the *Dnmt1* gene in adult mouse liver (38 ± 2.9%) [18]. We conclude that efficient adenine base editing can be achieved *in vivo* via single-AAV delivery of the Nme2-ABE8e system.

**Figure 6.**
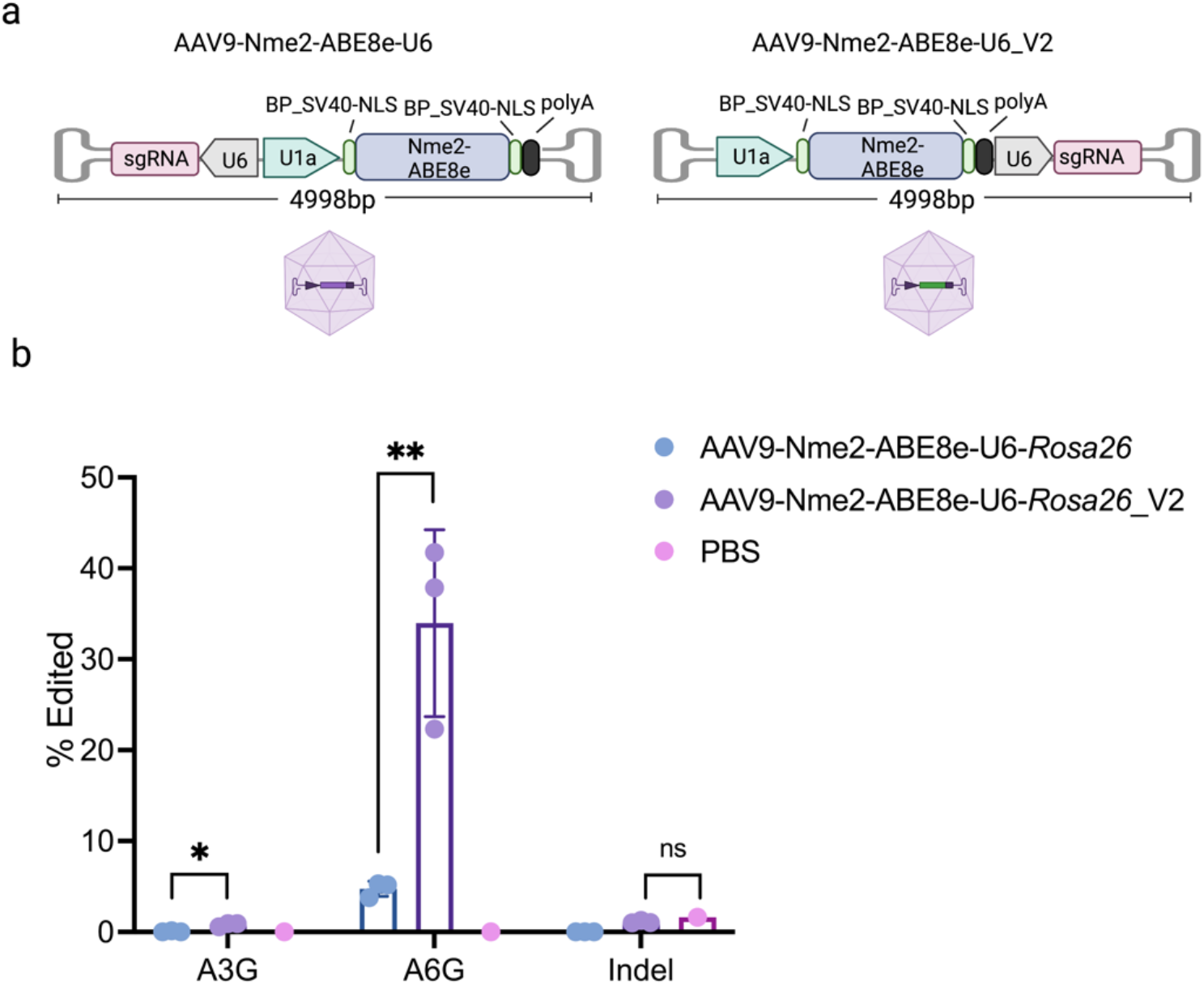
**a).** Schematic representation of the original (left) and the optimized AAV constructs (AAV9-Nm2-ABE8e-U6_V2, right). **b).** Quantification of the editing efficiency at the *Rosa26* locus by amplicon deep sequencing using mouse livers injected with indicated AAV (n = 3 mice in the AAV injected group, n = 1 mouse in the PBS injected group). Data from the original AAV configuration (**a**, left) is the same as that from the experiment shown in **Figure 5d**. Data represent mean ± SD, ns, P > 0.05; *, P<0.1; **, P < 0.01(one-way ANOVA).

## Discussion

Rapidly evolving and precise genome editing tools such as base editors and prime editors possess great potential to address the root causes of human genetic diseases [2,75,76]. However, the safe and effective delivery of genome editors remains a major challenge. Viral delivery using AAV, which is the only FDA-approved *in vivo* gene therapy vector to date, has a limited packaging capacity. Previous studies using AAV to deliver base editors or prime editors have been reported to be effective in rodents [7,8,10–13,15–18,21], but all of them required multiple vectors and most of them required high viral dosage [7,10–13,17,18]. To date, AAV administered at high dose has been reported to relate to severe toxicity or even death in nonhuman primates, piglets, and humans [23,24,77–79]. Engineering single-AAV deliverable genome editing tools has the potential to achieve therapeutic benefits at lower dosage, which would not only ease manufacturing burdens but also reduce likelihoods of serious adverse events [80,81].

Engineering new genome editors based on compact Cas9s is an alternative approach to address this issue. Previously, single-AAV delivery of Cas9 nucleases has been reported to generate NHEJ-based editing *in vivo* [26,30,82]. Precision editing via HDR has also been achieved with single-AAV systems, albeit with low efficiencies [27,83]. The only single-AAV BE system reported previously (which was not tested *in vivo*) was based on SauCas9, which has limited targeting range due to its PAM constraints. In this study, we constructed and characterized a compact, accurate adenine base editor, Nme2-ABE8e, which can target many sites that are inaccessible to SauCas9 base editors due to their distinct PAMs. We showed that Nme2-ABE8e with a sgRNA can be packaged into a single AAV vector, and a single intravenous injection in an adult disease mouse model of tyrosinemia reversed both the disease mutation and phenotypes. Furthermore, the dosage used in this study (4 x 10^11^ per mouse or 2 x 10^13^ vg/kg) is well below the 1 x 10^14^ vg/kg systemic doses that have been tolerated in clinical trials [77,78].

Although the editing efficiency was modest at the *Fah* disease locus, we reached therapeutic thresholds for HT1, and the initial editing efficiency of the AAV9-delivered Nme2-ABE8e-U6 construct exceeded that reported previously at this locus using other precision genome editing tools and delivery approaches [65–67,70]. Moreover, by optimizing transgene orientation in the AAV construct, we achieved significantly improved editing efficiency (~5-fold increase) by AAV9-delivered Nme2-ABE8e targeting the *Rosa26* gene, comparable to that achieved previously by dual-AAV delivered split-intein SpyCas9-ABEmax in adult mouse liver targeting another genomic site [18]. Future optimization of this system promises to further improve efficiency. There are multiple potential explanations for the inconsistent editing efficiencies we achieved with our first-generation Nme2Cas9 ABE that suggest directions for future improvement. First, structural analyses of Nme2Cas9 [84] indicate that the position of the N-terminally fused TadA8e domain relative to the predicted path of the displaced strand is not optimal. With other effectors, domain-inlaid deaminase fusions have proven to be advantageous in some contexts [29,54], and the same is likely to be true with Nme2-ABEs. Second, the current Nme2-ABE8e has a wide editing window that could result in increased bystander editing. For example, we observed higher editing at a bystander adenine (A16) at the *Fah* locus compared to the on-target adenine at A13. Optimizations of linker composition and length [3,53], as well as domain-inlaid systems [29,54–58], may confer greater control over the editing window and improved efficiency in editing the intended position.

Our studies also suggest factors that must be considered for optimal guide expression. In contrast to a previous study that showed no significant difference between the U6 and miniU6 promoters in supporting Cas9 editing efficiency [62], we found that the construct with the miniU6 promoter was consistently less efficient than that with the U6 promoter, including via AAV delivery. Because the previous study compared these promoters in T cells by lentivirus transduction, our observations may only apply to certain delivery strategies and cellular contexts.

In summary, we have engineered and characterized Nme2-ABE8e editing in mammalian cell culture and achieved efficient *in vivo* editing by delivery of a single AAV vector. To our knowledge, this is the first single-AAV-delivered *in vivo* base editing reported to date [29]. We anticipate that Nme2-ABE8e, with its distinct PAM specificity, editing window, and high accuracy, will provide additional targetability, safety, and therapeutic potential for genome engineering applications.

## Methods

### Cell culture

HEK293T cells (ATCC CRL-3216), ABE reporter cells, MEF cells, and mouse N2a cells (ATCC CCL-131) were cultured in in Dulbecco’s Modified Eagle Media (DMEM, Genesee Scientific Cat. #: 25-500) supplemented with 10% Fetal Bovine Serum (Gibco Cat. #: 26140079). Rett syndrome human patient-derived fibroblasts (hPDFs) were obtained from the Rett Syndrome Research Trust and cultured with DMEM supplemented with 15% FBS and 1x non-essential amino acids (Gibco Cat. #: 11140050). All cells were incubated in a 37°C incubator with 5% CO_2_.

### Molecular cloning

To generate the CMV-Nme2-ABE8e and the CMV-Nme2-ABE7.10 plasmids used in **Figure 1**, the Nme2-ABE8e, Nme2-ABE7.10, Spy-ABE8e, and Spy-ABE7.10 constructs were cloned into the pCMV-PE2 vector backbone (Addgene #132775) by Gibson assembly. Briefly, the pCMV-PE2 plasmid was digested with NotI and PmeI restriction enzymes, and the plasmid backbone was then Gibson-assembled with five fragments: N-terminal NLS, TadA8e (for Nme2-ABE8e or Spy-ABE8e), or TadA-TadA*7.10 (for Nme2-ABE7.10 or Spy-ABE7.10), the linker, Nme2Cas9-D16A or SpCas9-D10A nickase, and the C-terminal NLS. The ABE reporter construct was cloned by site-directed mutagenesis to change the 47^th^ amino acid (Glutamine, CAG) of the mCherry coding sequence to a stop codon (TAG). The ABE reporter was further cloned into a lentiviral transfer vector backbone (Addgene #99373) by Gibson assembly. The sgRNA expression plasmids used in **Figure 1** were cloned from pBluescriptSKII (Addene #74705) in two steps. First, pBluescriptSKII was digested by NotI and XbaI, and a gene fragment that contains a U6 promoter, a type-IIS restriction cloning site (BfuAI), and a tracrRNA was assembled into the backbone by Gibson assembly. The plasmids were further digested with BfuAI and ligated to the annealed oligos to insert the guide sequences. The single-AAV vector plasmids in **Figure 3a** were cloned from the Addgene #119924 plasmid by replacing the Nme2Cas9 sequence with the Nme2-ABE8e sequence, and then subsequently re-cloned to encode different NLS configurations and the miniU6 promoter by restriction enzyme digestion and Gibson assembly. The NLS configuration of Nme2-ABE8e mRNA used in **Figure 2b** was 2x_BPSV40, and the plasmid used in **Figure 2d** was the single-AAV Nme2-ABE8e-U6 plasmid shown in **Figure 3a** (2xBP_SV40). To clone the AAV-Nme2-ABE8e_V2 plasmid in **Figure 6**, first, the 2xBP_SV40 plasmid from **Figure 3a** was digested by PmeI and NotI, the AAV backbone fragment was then Gibson-assembled with the fragment containing U1a-Nme2-ABE8e, and the fragment containing U6-sgRNA with homology sequences by overhang PCR. Most of the plasmids used in **Supplementary Figure 3a** were previously deposited in Addgene (AcrIIC3, Addgene #85713; AcrIIC4, Addgene #113434; Acr-E2, Addgene #85677). To clone the plasmid expressing AcrIIA4, Addgene #85713 plasmid was digested with XhoI and BamHI and Gibson-assembled with a gene fragment containing AcrIIA4 coding sequence [46]. In **Supplementary Figure 3b**, the AcrIIC3-MRE122 plasmid was cloned from Addgene plasmid #129531 by replacing the *Rosa26*-targeting guide with the DS12-targeting guide through restriction enzyme cloning. Subsequent replacement of the coding sequence of AcrIIC3 with AcrIIC4 generated the AcrIIC4-MRE122 plasmid. Sequences of plasmids first described in this paper can be found in the **Supplementary note** and will be made available from Addgene.

### ABE reporter HEK293T cell line

Lentivirus was produced following instructions from Addgene (https://www.addgene.org/protocols/lentivirus-production/). Briefly, HEK293T cells were transfected with the transfer vector and the packaging plasmids psPAX2 (Addgene #12260) and pMD2.G (Addgene #12259), using Lipofectamine 3000 (ThermoFisher Cat. #: L3000015). Two days later, the medium was collected and filtered through a 0.45 μm filter (Cytiva Cat. #: 6780-25040) to remove cell debris. The viral titer was determined using Lenti-X™GoStix™ (Takara Bio Cat. #: 631280). HEK293T cells were transduced with lentivirus encoding the ABE reporter at varying dilutions (1:10, 1:100, 1:500, and 1:1000) in the presence of 8 μg/ml polybrene (Millipore Sigma Cat. #: TR-1003-G). Three days after transduction, the media was removed, and fresh media was supplemented with 2 μg/ml Puromycin to select cells expressing the full-length reporter construct. Seven days after selection, the puromycin-resistant cells were collected. Single-cell clones were established by serial dilution in 96-well plates.

### Fluorescent reporter assay

Forty-eight hours after transfection, cells were trypsinized and harvested into microcentrifuge tubes. After centrifuging at 300 x g for 3 mins, cells were resuspended into 150ul 1xPBS for flow cytometry analysis (MACSQuant VYB). For each sample, 10,000 events were counted for FACS analysis. Data was analyzed using Flowjo v10. Representative gating strategy can be found in **Supplementary Figure 2**.

### *In vitro* transcription of Nme2-ABE8e mRNA

Nme2-ABE8e mRNA was *in vitro* transcribed using the NEB HiScribe T7 RNA synthesis kit, from 500ng of a linearized template. Uridine was fully substituted with 1-methylpseudouridine, and mRNA was capped co-transcriptionally using CleanCap AG analog (TriLink Biotechnologies Cat. #N-1081 and N-7113, respectively). All enzymes were purchased from New England Biolabs. Transcription was conducted according to the manufacturer’s protocol with the following amendments: transcriptions were completed in 1 × NEB HiScribe transcription buffer, and 4 mM CleanCap AG was used during transcription. Transcription reactions were incubated at 37 °C for 2 hours then treated with 0.4 U/μl DNase I (final concentration) for 15 min at 37 °C. mRNAs were purified with NEB Monarch RNA purification columns and treated with pyrophosphatase for 1 hour with 0.25 U/μg Antarctic phosphatase (final concentration) in 1X Antarctic phosphatase buffer. The final product was further purified with a NEB Monarch RNA column and eluted in water.

### Transfection and electroporation

For plasmid transfection, cells were seeded in 24-well plates at 80,000 cells per well in culture media and incubated overnight. Briefly, plasmids were transfected at 400ng per well when targeting the ABE reporter, and 1ug per well for the endogenous target sites. Specifically, in **Figure 1c**, **1e**, **2d**, **3b**, **3c**, and **Supplementary Figure 3**, plasmids were transfected using Lipofectamine3000 (ThermoFisher Cat #L3000001), while in **Figure 1d** and **Supplementary Figure 2**, plasmids were transfected using Lipofectamine2000 (ThermoFisher Cat #11668030), following the manufacturer’s protocols. Electroporation was performed using Neon TransfectionSystem 10 ul kit (ThermoFisher Cat #: MPK 1096), with the following electroporation parameters: Pulse voltage (1650 v), Pulse width (20 ms), Pulse number (1). Specifically, in **Figure 2a**, 264 ng Nme2-ABE8e mRNA and 100 pmole of sgRNA were electroporated into 50,000 Rett Syndrome hPDFs, while in **Figure 4c**, 1 ug plasmid DNA was electroporated into 100,000 MEF cells.

### AAV production

AAV vector packaging was done at the Viral Vector Core of the Horae Gene Therapy Center at the UMass Chan Medical School as previously described [86]. Constructs were packaged in AAV9 capsids and viral titers were determined by digital droplet PCR and gel electrophoresis followed by silver staining.

### AAV genomic DNA extraction and alkaline agarose gel electrophoresis

Genomic DNAs were extracted from 10^11^ vg AAV by incubating with 20 units of DNase I (ThermoFisher Cat. #: EN0521) at 37 °C for 30 minutes and then with an equal volume of 2 x Pronase solution (Sigma-Aldrich, Cat. # 10165921001) for 4 hours. The genomic DNA was subsequently purified by phenol-chloroform (ThermoFisher Cat. #:15593-049) extraction and ethanol precipitation. DNA pellets were resuspended in water and analyzed by alkaline agarose gel electrophoresis and SYBR Gold staining.

### Mouse experiments

All animal study protocols were approved by the Institutional Animal Care and Use Committee (IACUC) at UMass Chan Medical School. The *Fah*^PM/PM^ mice were kept on water supplemented with 10 mg/L 2-(2-nitro-4-trifluoromethylbenzoyl)-1,3-cyclohexanedione (NTBC; Sigma-Aldrich, Cat. #: PHR1731-1G). Mice with more than 20% weight loss were humanely euthanized according to IACUC guidelines. For hydrodynamic tail vein injections, plasmids were prepared by EndoFree Plasmid Maxi Kit (Qiagen Cat. #12362). Briefly, 30 ug SpCas8-RA6.3 and 30 ug sgRNA plasmids, or 60 ug Nme2-ABE8e single AAV plasmids were suspended in 2 ml saline and injected via tail vein within 5-7 seconds into 10-week-old *Fah*^PM/PM^ mice. Mice were euthanized 7 days after injection and livers were collected for analysis. For AAV injection, a dosage of 4 x 10^11^ vg per mouse (in 200 ul saline) was tail-vein injected into 8-week-old *Fah*^PM/PM^ mice.

### Genomic DNA extraction from cultured cells and mouse liver

For cultured cells, genomic DNAs were extracted 72 hours after plasmid transfection, or 48 hours after mRNA and sgRNA electroporation. Briefly, cell culture media were aspirated, and cells were washed with 1x PBS. Genomic DNAs were prepared using QuickExtract DNA Extraction Solution (Lucigen) following the manufacturer’s protocols. To extract genomic DNA from mouse livers, all 5 lobes of mouse liver were combined and pulverized in liquid nitrogen, and 15 mg of the tissue from each mouse liver was used for genomic DNA extraction using GenElute Mammalian Genomic DNA Miniprep Kit (Millipore Sigma Cat. #: G1N350).

### Amplicon deep sequencing and data analysis

Genomic DNA was amplified by PCR using Q5 High-Fidelity 2X Master Mix (NEB Cat. #: M0492) for 20 cycles. One microliter of the unpurified PCR product was used as a template for 20 cycles of barcoding PCR. The barcoding PCR reactions were further pooled and gel-extracted using Zymo gel extraction kit and DNA clean & concentrator (Zymo research Cat. #: 11-301 and 11-303) and quantified by Qubit 1x dsDNA HS assay kits (Thermo Fisher Scientific, Cat #: Q32851). Sequencing of the pooled amplicons was performed using an Illumina MiniSeq system (300-cycles, FC-420-1004) following the manufacturer’s protocol. The raw MiniSeq output was de-multiplexed using bcl2fastq2 (Illumina, version 2.20.0) with the flag --barcode-mismatches 0. To align the generated fastq files and to quantify editing efficiency, CRISPResso2 [87] (version 2.0.40) was used in batch mode with base editor output and the following flags: -w 15, -q 30. Indel frequency = (insertions reads + deletions reads)/all aligned reads x 100%.

### Immunohistochemistry (IHC)

Mice were euthanized by CO_2_ asphyxiation and livers were fixed with 10% neutral buffered formalin (Epredia Cat. #: 5735), sectioned at 10um, and stained with hematoxylin and eosin for pathology analysis. For IHC, liver sections were dewaxed, rehydrated, and stained using an anti-FAH antibody (Abcam Cat. #: ab83770) at 1:400 dilution as described previously [67].

### Reverse transcription PCR

All 5 lobes of each mouse liver were combined and pulverized in liquid nitrogen. Total RNA was extracted from 50 mg of liver tissue using TRIzol reagent (ThermoFisher Cat. #: 15596026) and reverse-transcribed using SuperScript III First-Strand Synthesis System (ThermoFisher Cat. #:18080051). PCR was performed using primers previously described [67].

## Author contributions

H.Z., X.D.G., and E.J.S conceived the study. H.Z. designed, performed, and analyzed the *in vivo* experiments. H.Z., N.B., O.O., P.L., and X.D.G., designed, performed, and analyzed the *in vitro* experiments. T.R. and H.C. analyzed the deep sequencing data. E.J.S., W.X., and S.A.W. supervised research. H.Z. and E.J.S. wrote the manuscript with contributions from N.B., P.L., and H.C. All authors edited the manuscript.

## Acknowledgements

We thank Yueying Cao and Greg Cottle for their assistance with the mouse colony maintenance and tail vein injections. We thank Zexiang Chen for his help with the *in vitro* mRNA transcription, Nadia Amrani for providing the AcrIIA4 plasmid, Tingting Jiang for providing *Fah*^PM/PM^ MEF cells and Spy-RA6.3 plasmid, and NamKung Suk for his assistance with the alkaline gel electrophoresis. We also thank the UMMS Viral Vector Core for AAV packaging service, the UMMS Morphology Core for tissue sectioning and IHC staining, and the Rett Syndrome Research Trust for patient-derived fibroblasts. We are grateful to all members of the Gao, Wolfe, Xue and Sontheimer labs for their valuable discussions, advice, and helpful feedback.

---

**Supplementary Figure 1.**
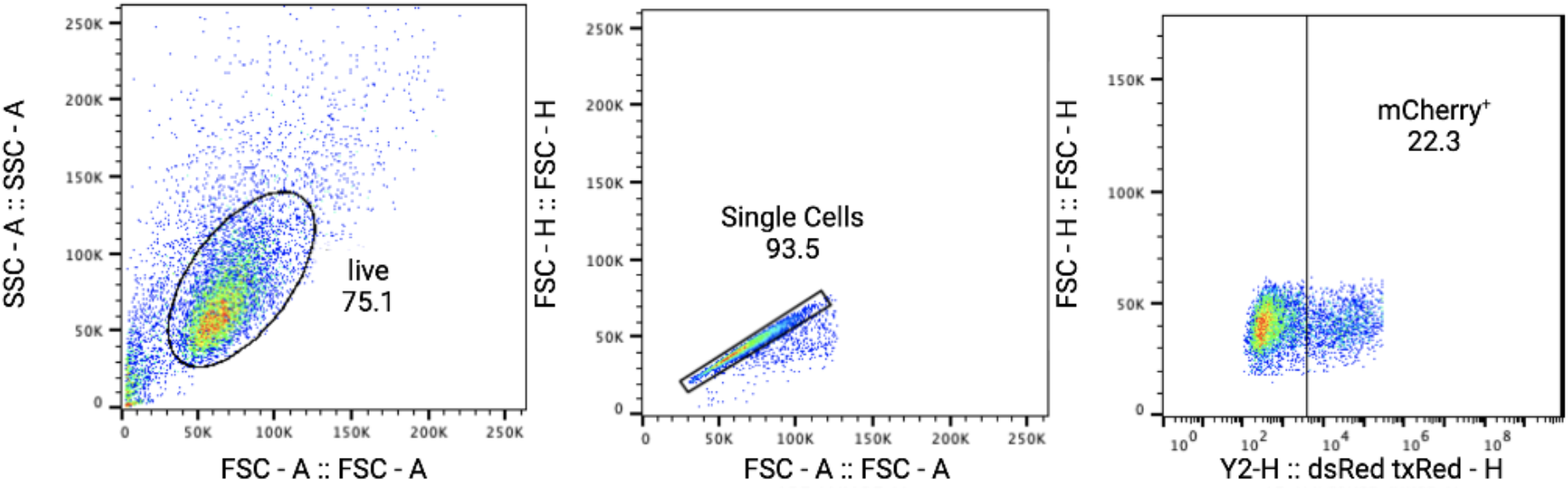
Representative flow cytometry gating strategy for the ABE reporter cell line.

**Supplementary Figure 2.**
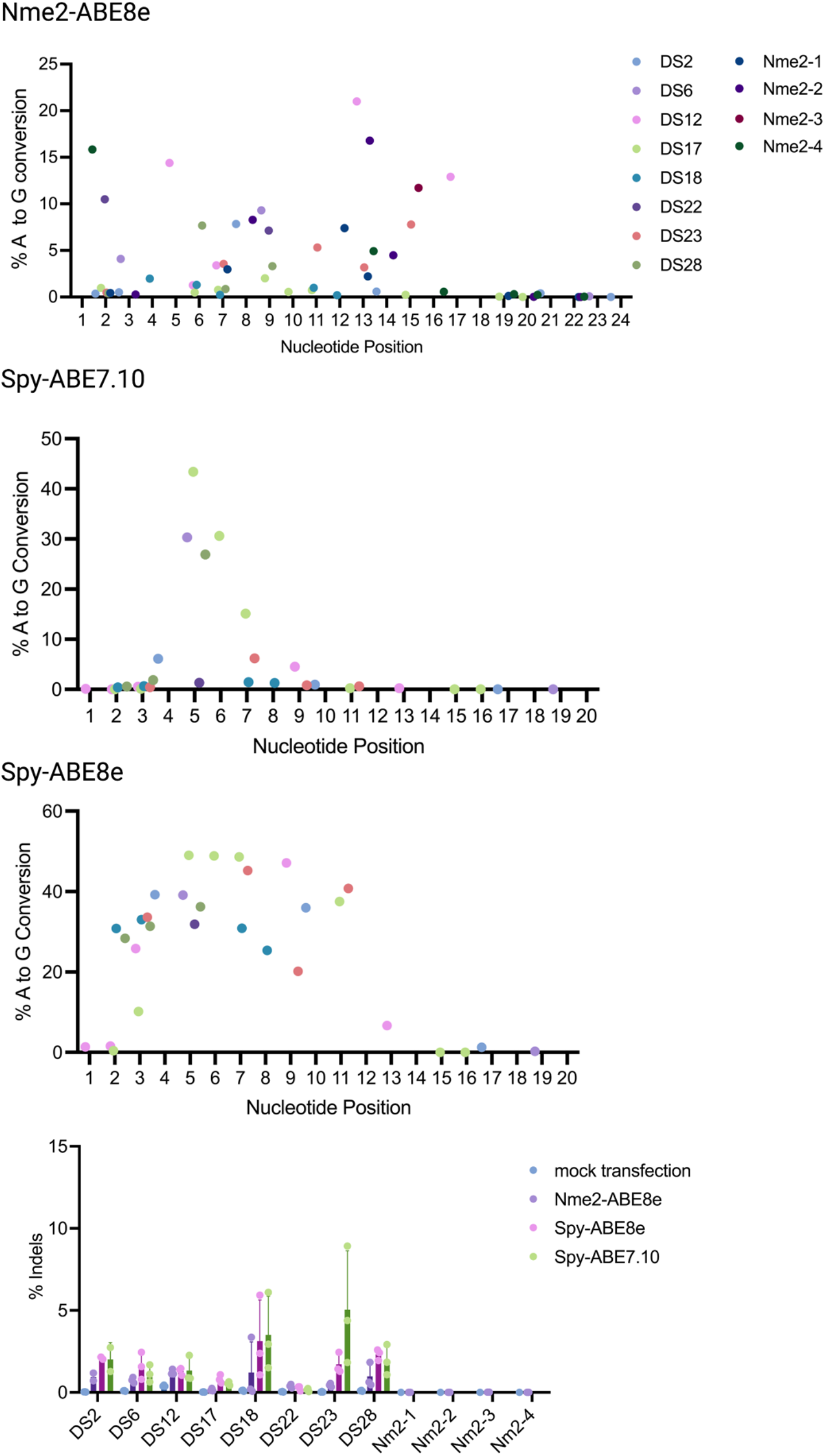
Summary of the individual A-to-G conversion efficiency and the indel efficiency (bottom) at 12 target sites for Nme2-ABE8e, which includes 8 dual-target sites (DS 2 - 28) and 4 Nme2Cas9-specific target sites (Nme2 1 - 4), and 8 dual-target sites for Spy-ABE7.10 and Spy-ABE8e. Each data point represents the A-to-G conversion efficiency at the indicated nucleotide position measured by amplicon deep sequencing (n = 3 biological replicates).

**Supplementary Figure 3.**
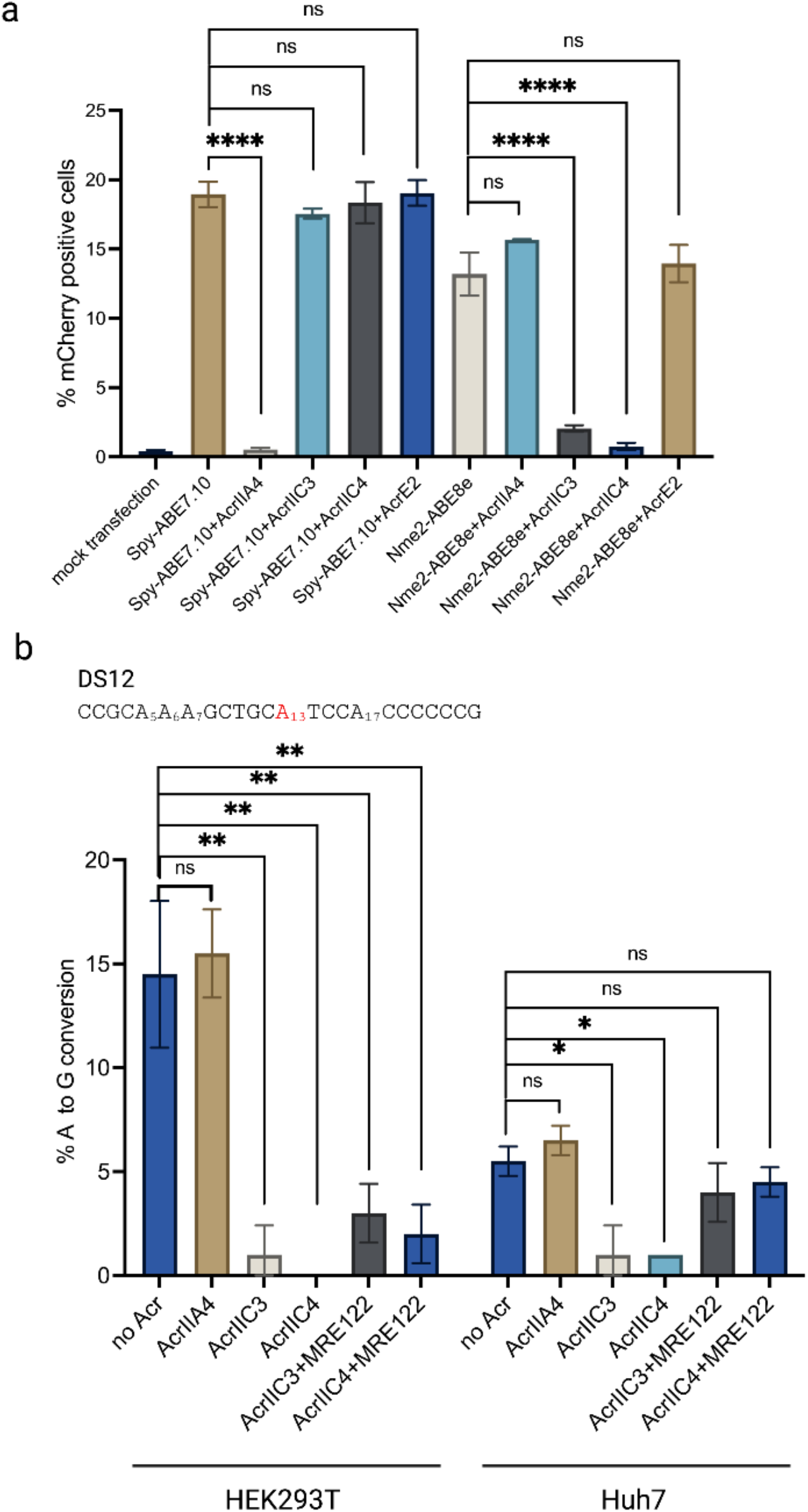
**a).** Anti-CRISPR proteins inhibit the activity of either Spy-ABE7.10 or Nme2-ABE8e in the ABE reporter cell line as a function of their previously defined nuclease specificity (n = 2 biological replicates). **b).** Editing efficiency at the A12 (the highest edited adenine within the target) of the DS12 site by Nme2-ABE8e controlled by microRNA-repressible anti-CRIPSR proteins. Data generated by Sanger sequencing and EditR analysis (n = 2 biological replicates). Data represent mean ± SD; ns, P > 0.05; *, P < 0.05; **, P < 0.01; ****, P < 0.0001 (one-way ANOVA).

**Supplementary Figure 4.**
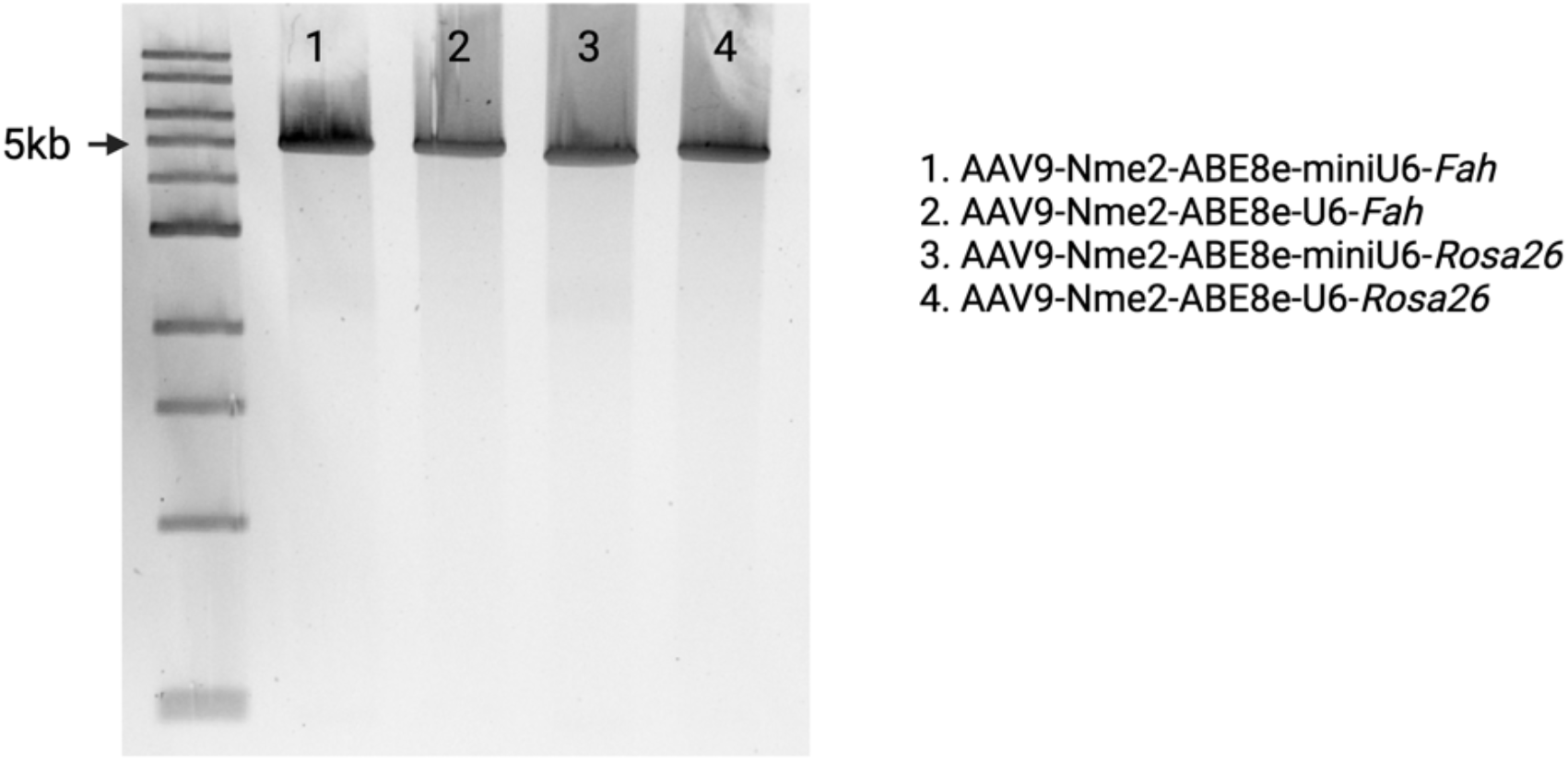
Alkaline gel electrophoresis of AAV9 genomic DNA.

**Supplementary Figure 5.**
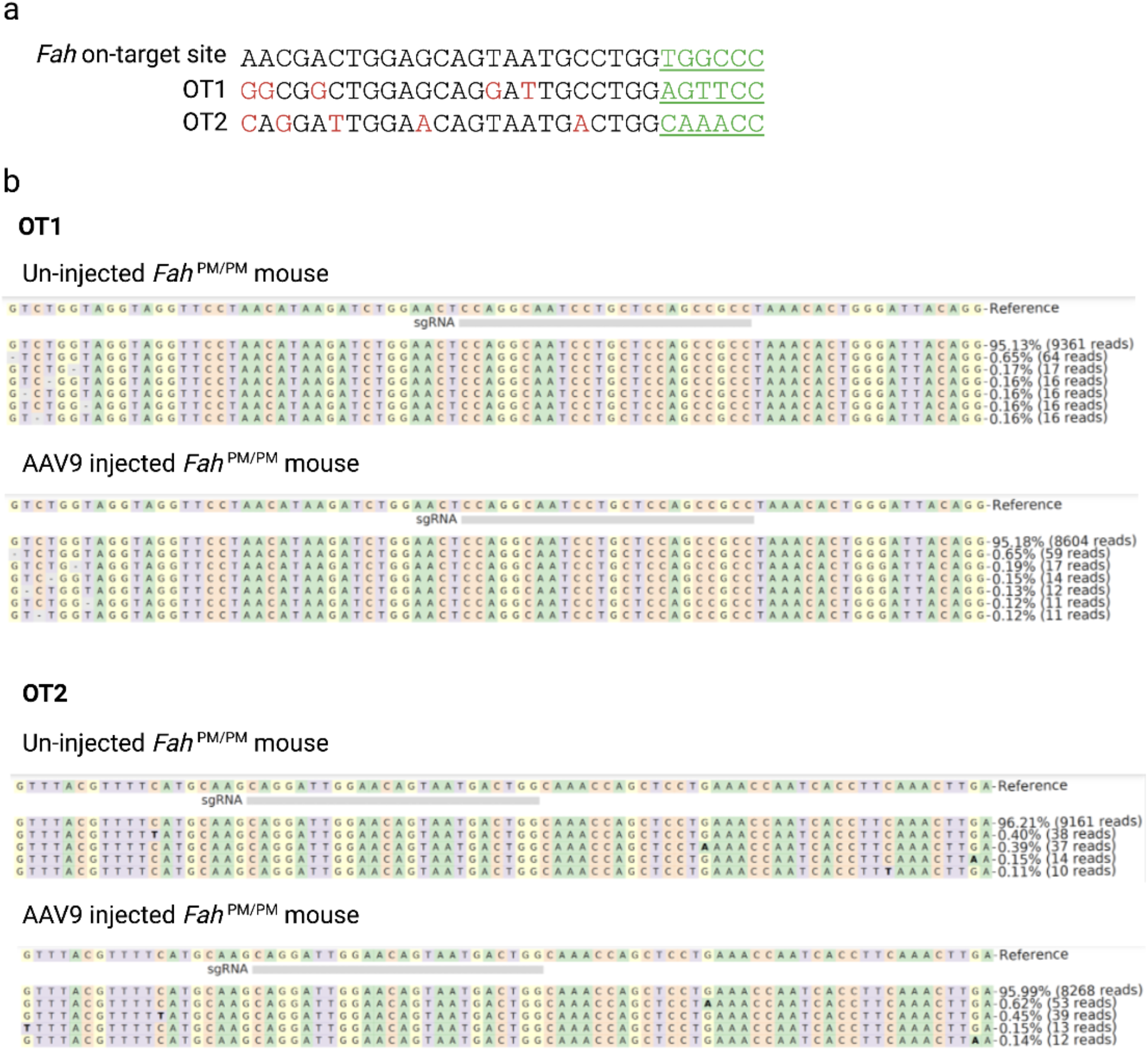
**a).** Sequence of the *Fah* on-target site and two top-rated Cas-OFFinder predicted off-target sites for Nme2-ABE8e. Bases that are different from the on-target site are labeled in red. PAM, green, underlined. **b).** Representative amplicon deep sequencing reads at the predicted off-target sites in mice injected with AAV9 expressing Nme2-ABE8e and sgRNA-*Fah*.

## Supplementary note

Nucleotide sequences of plasmids

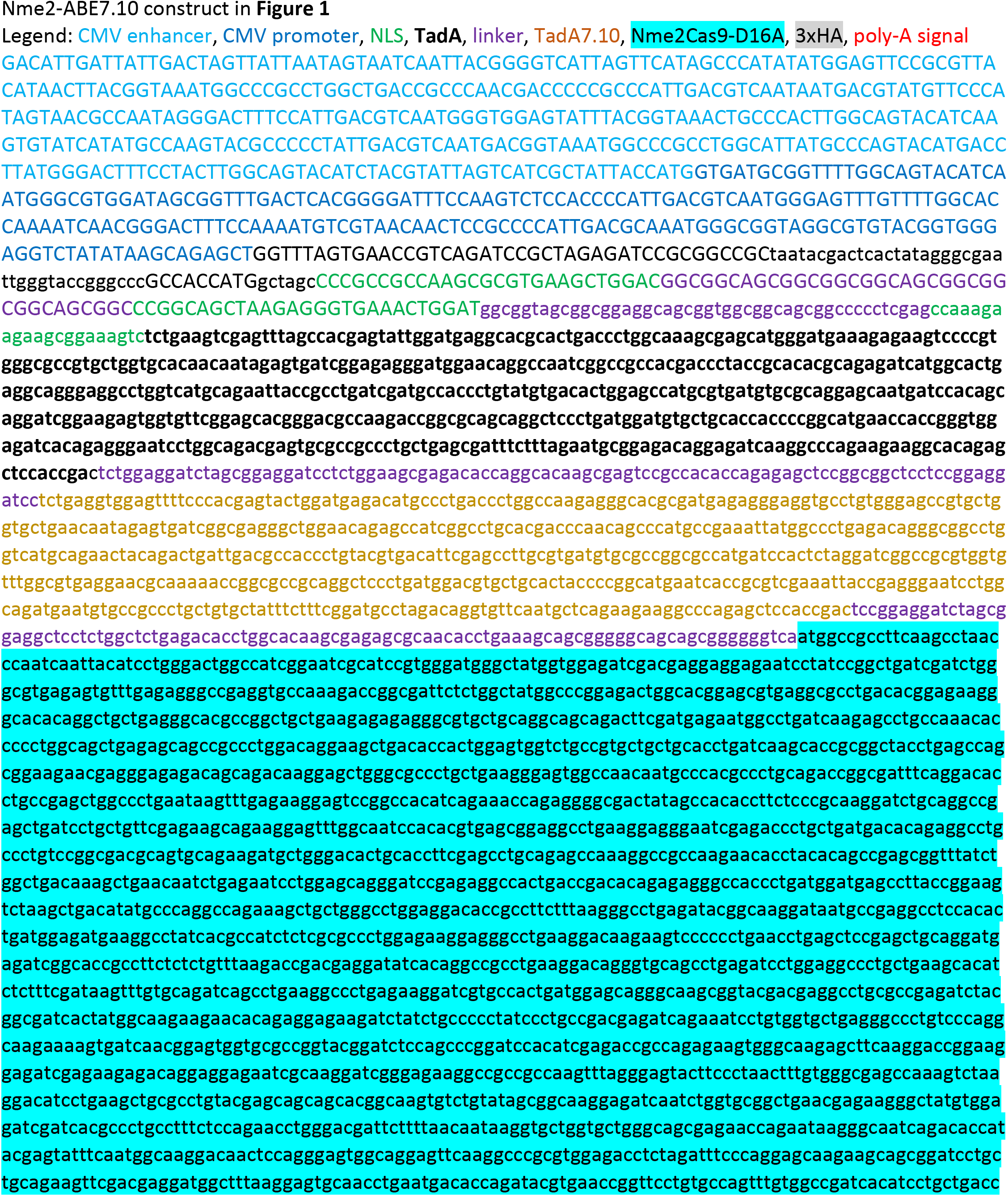

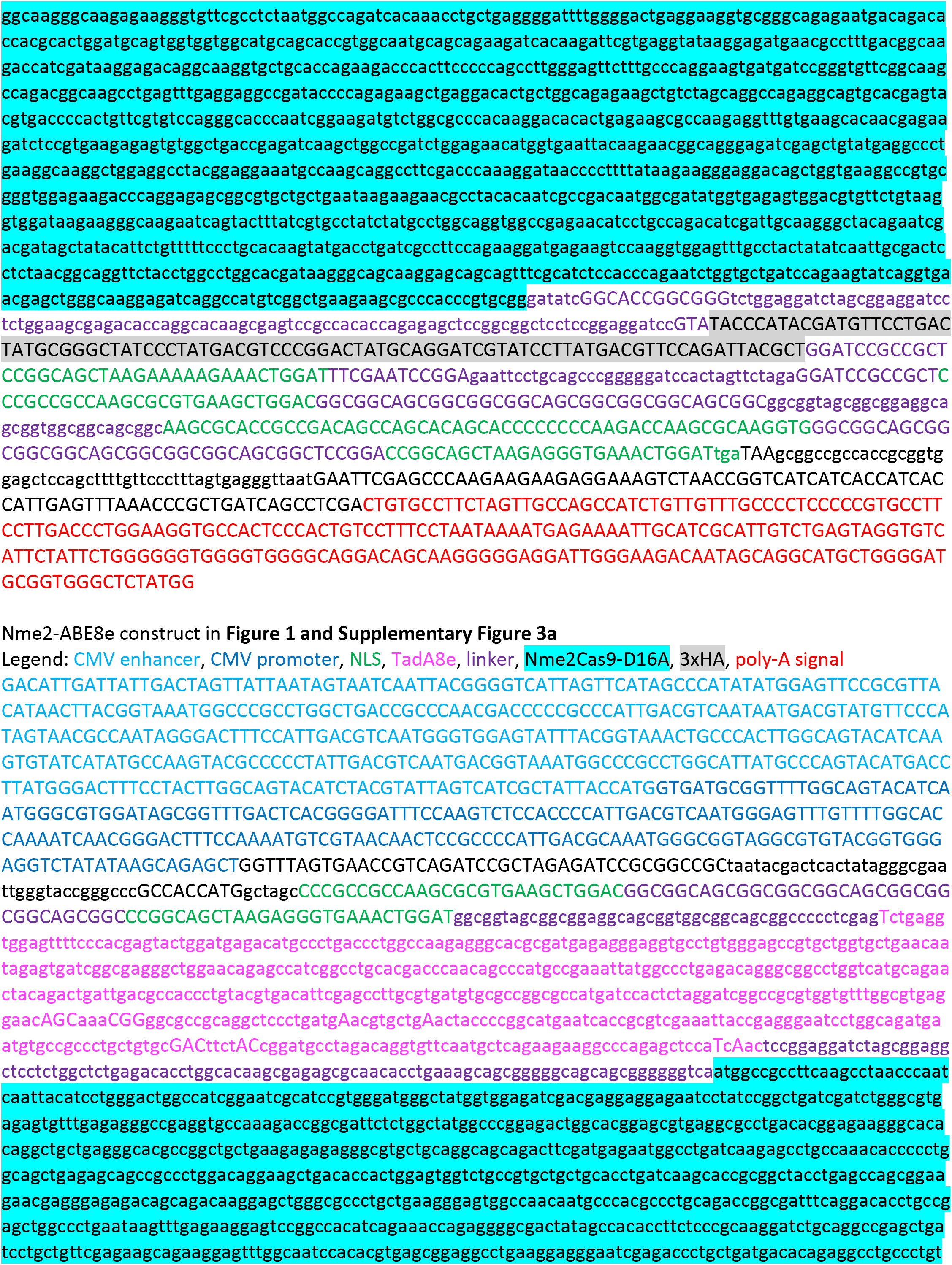

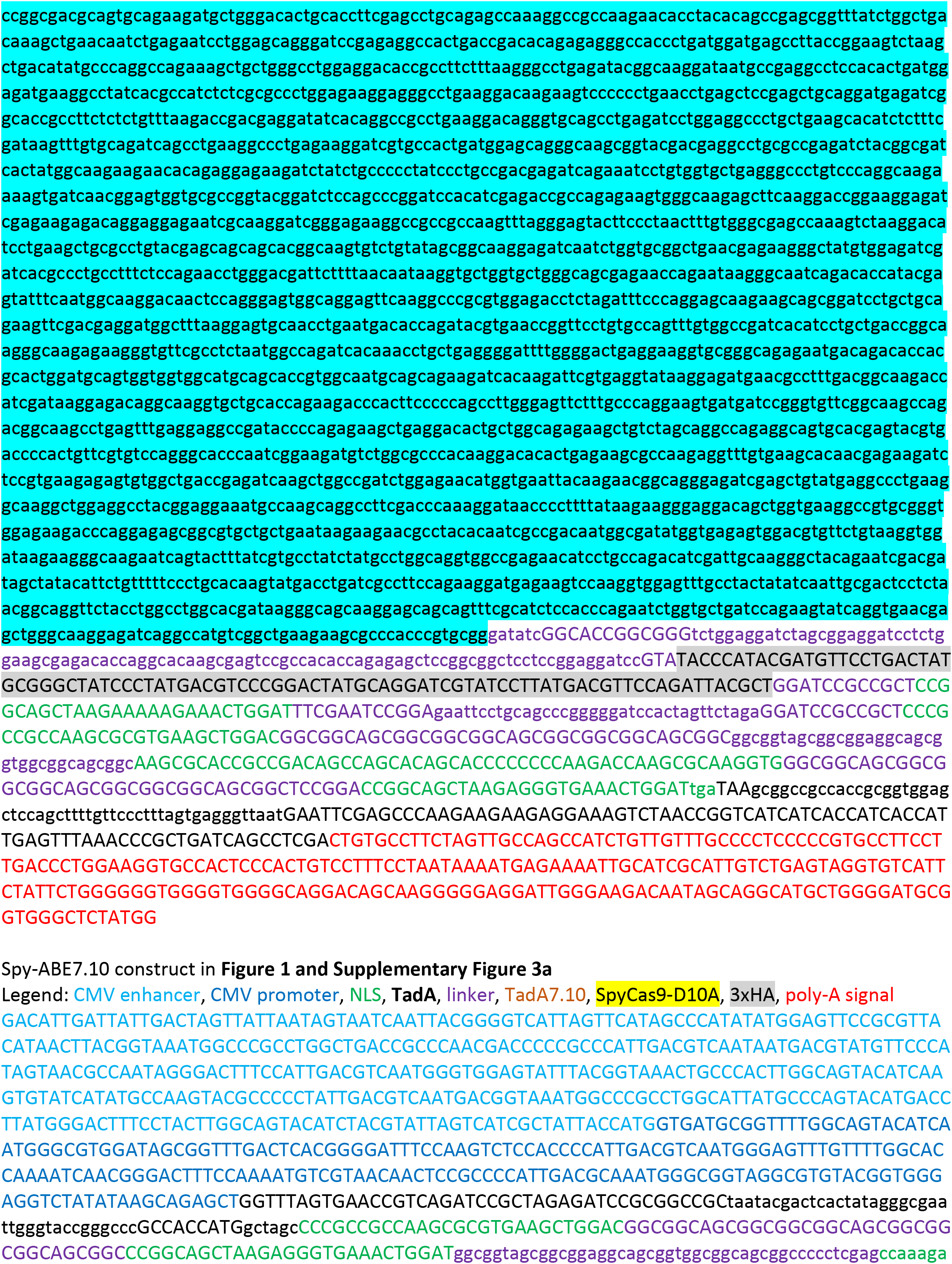

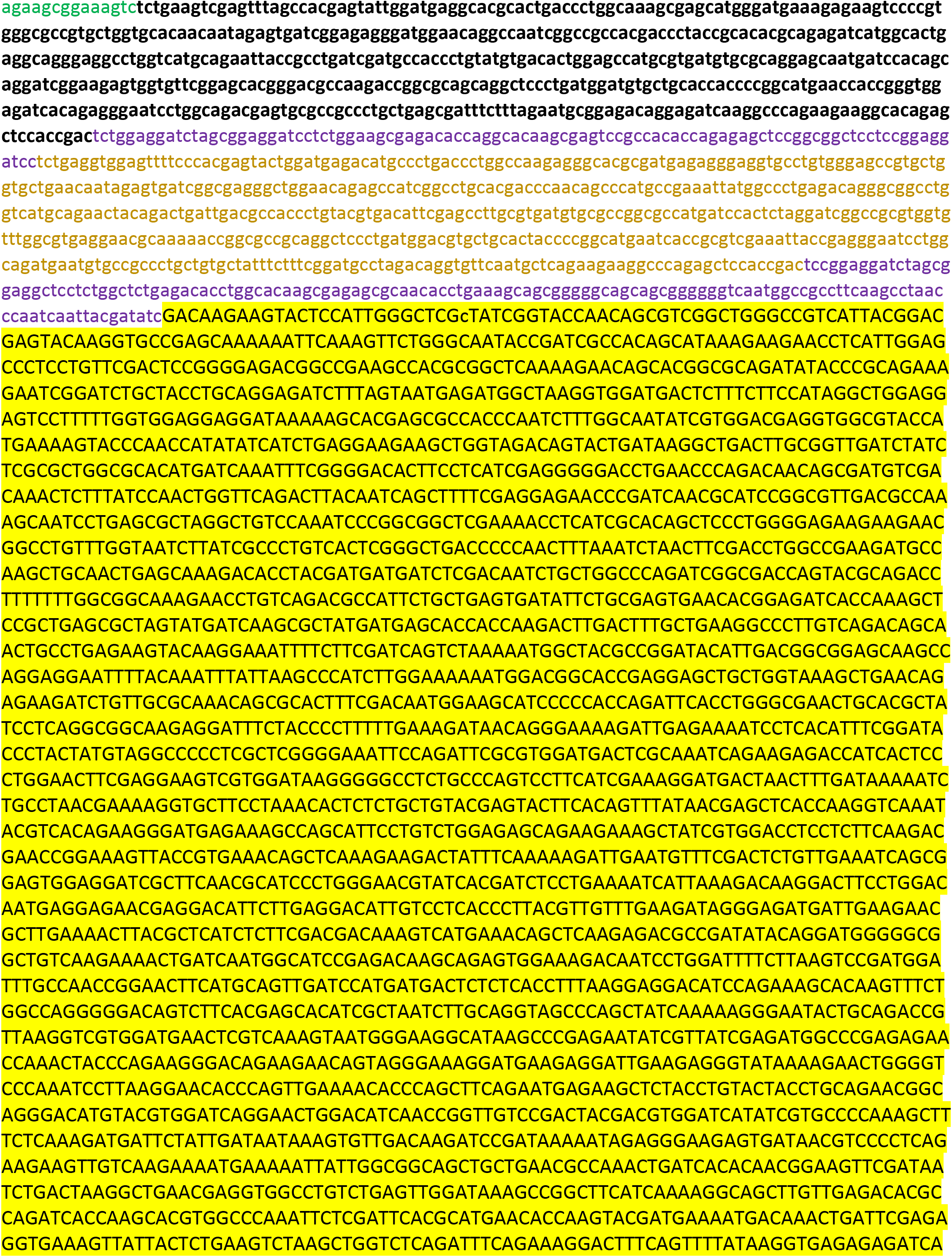

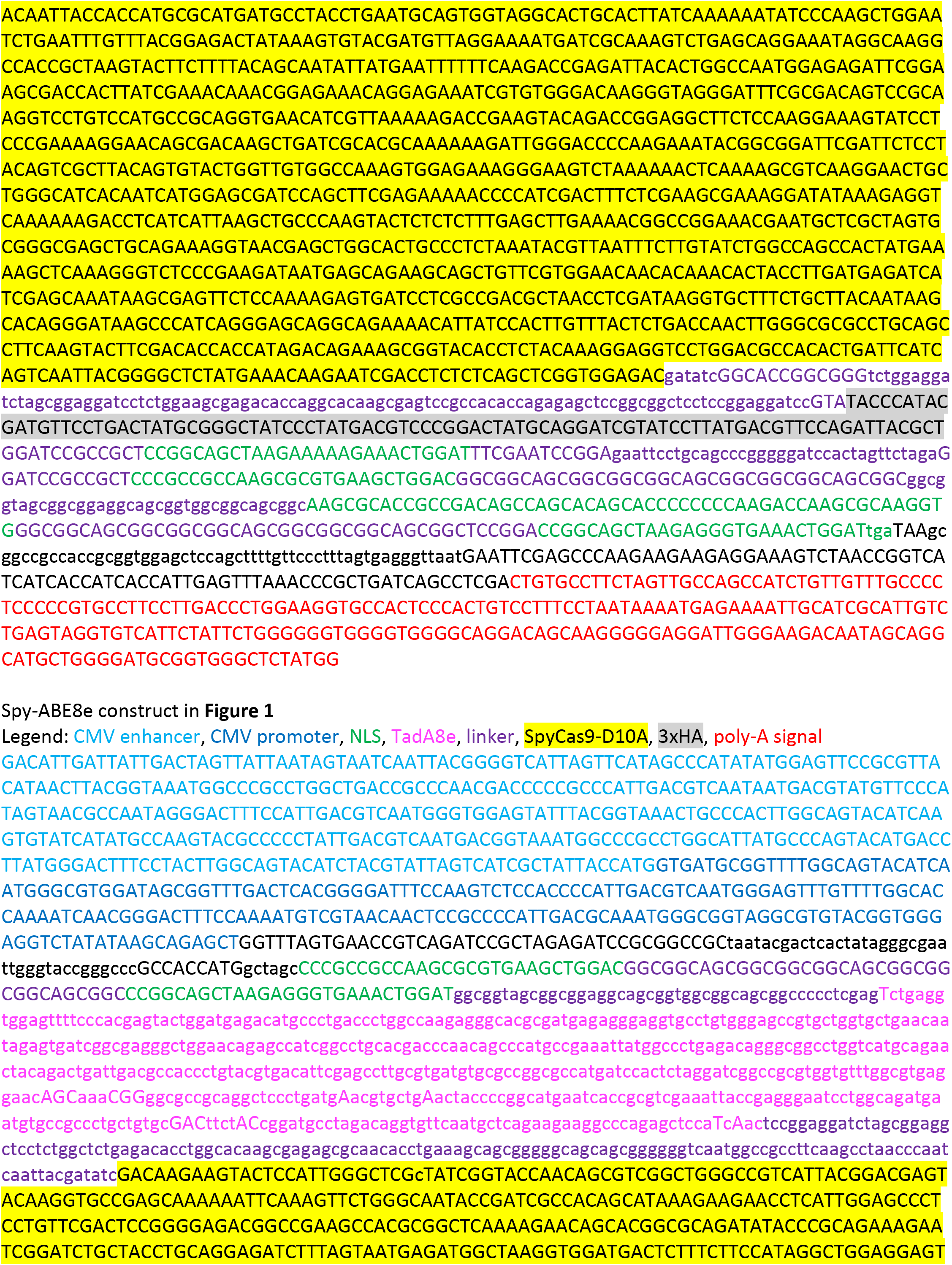

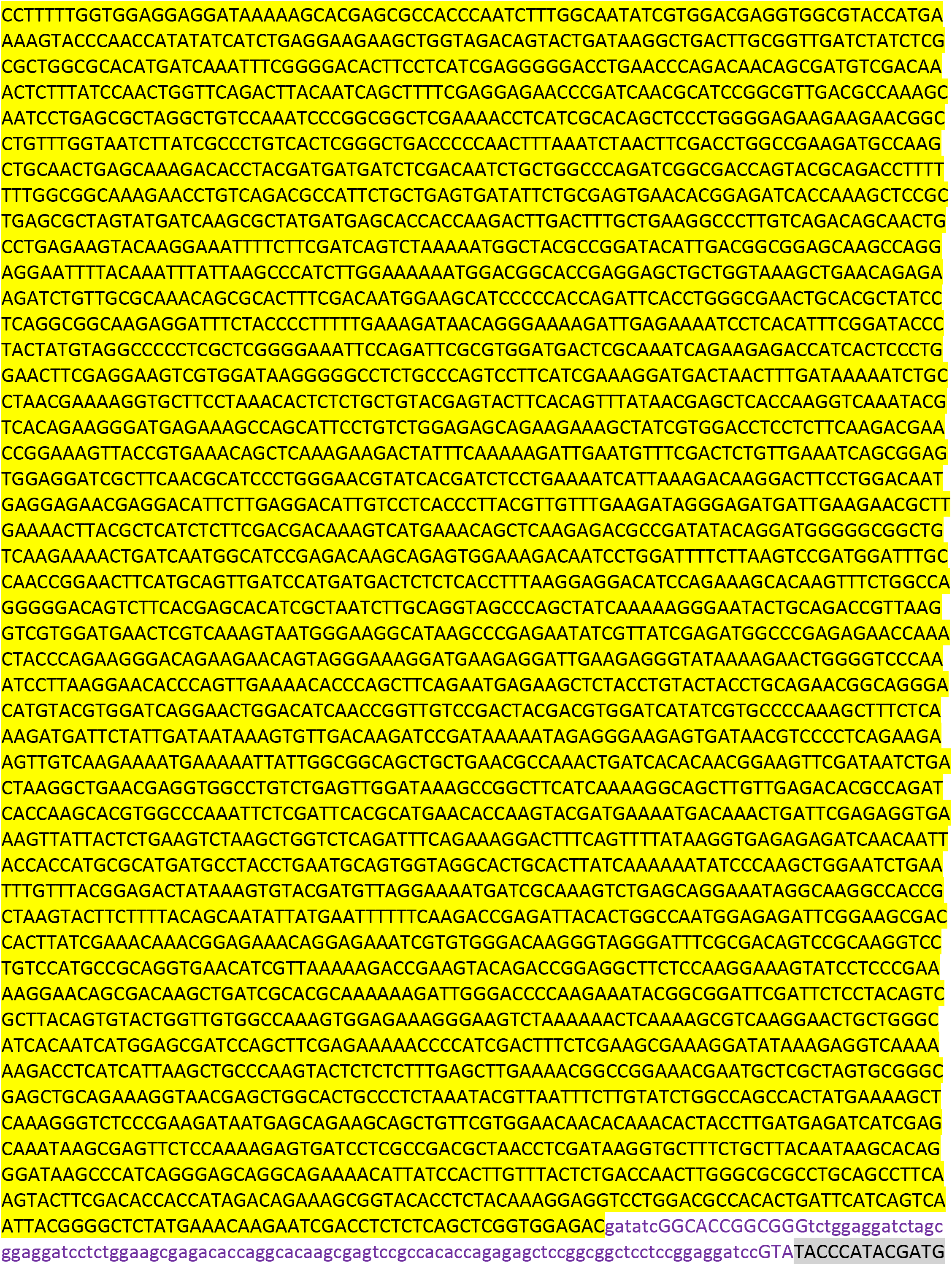

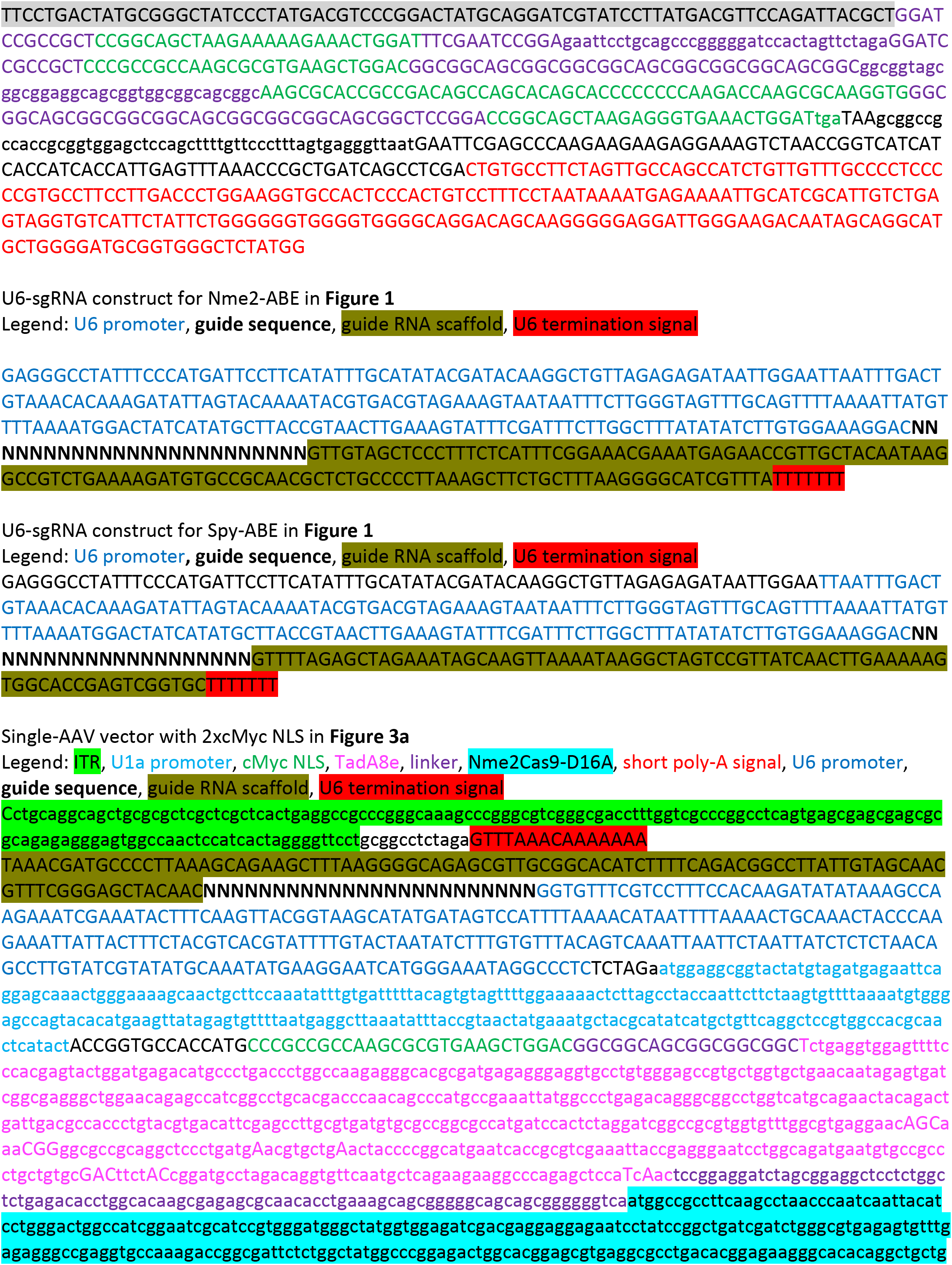

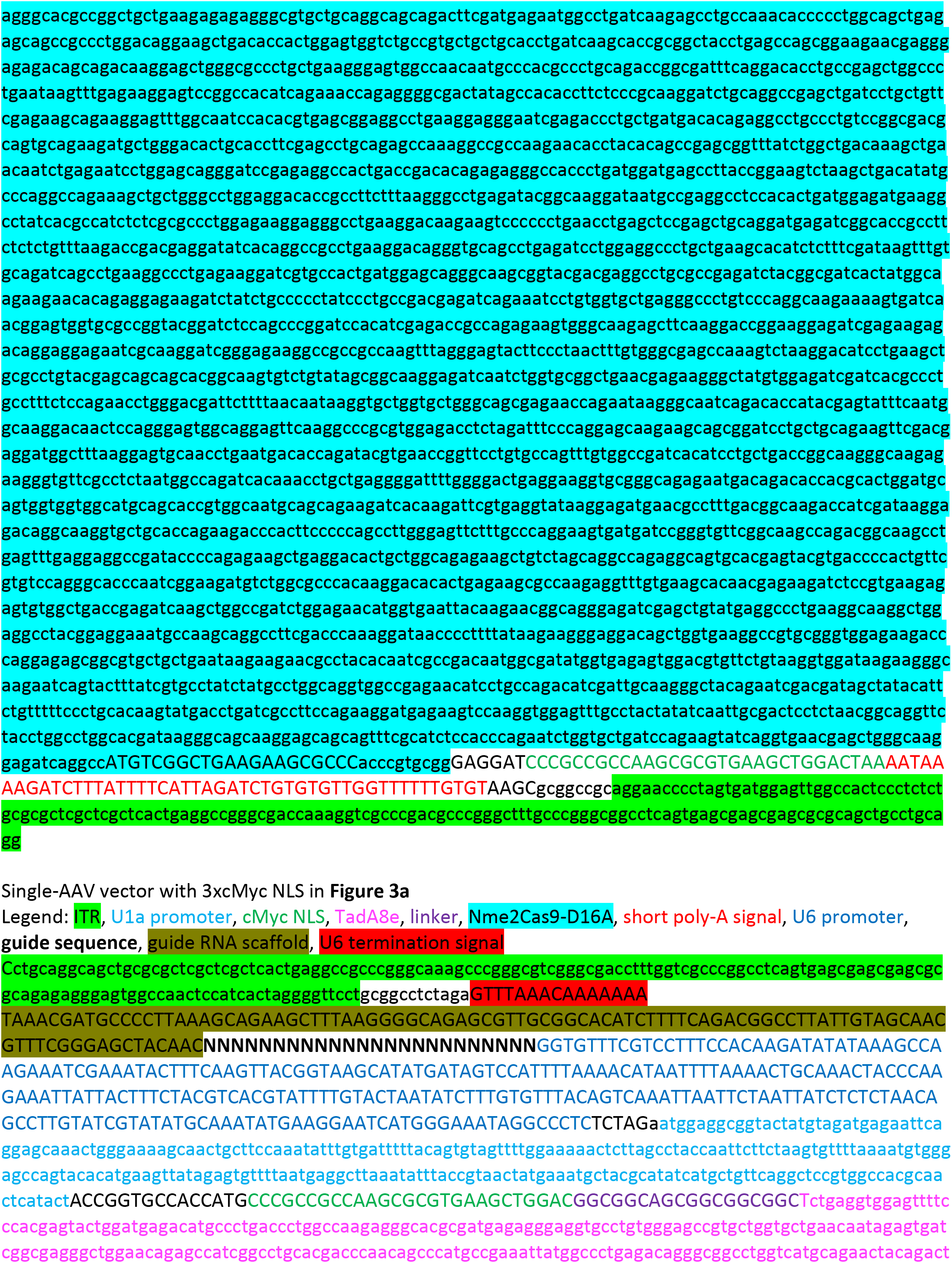

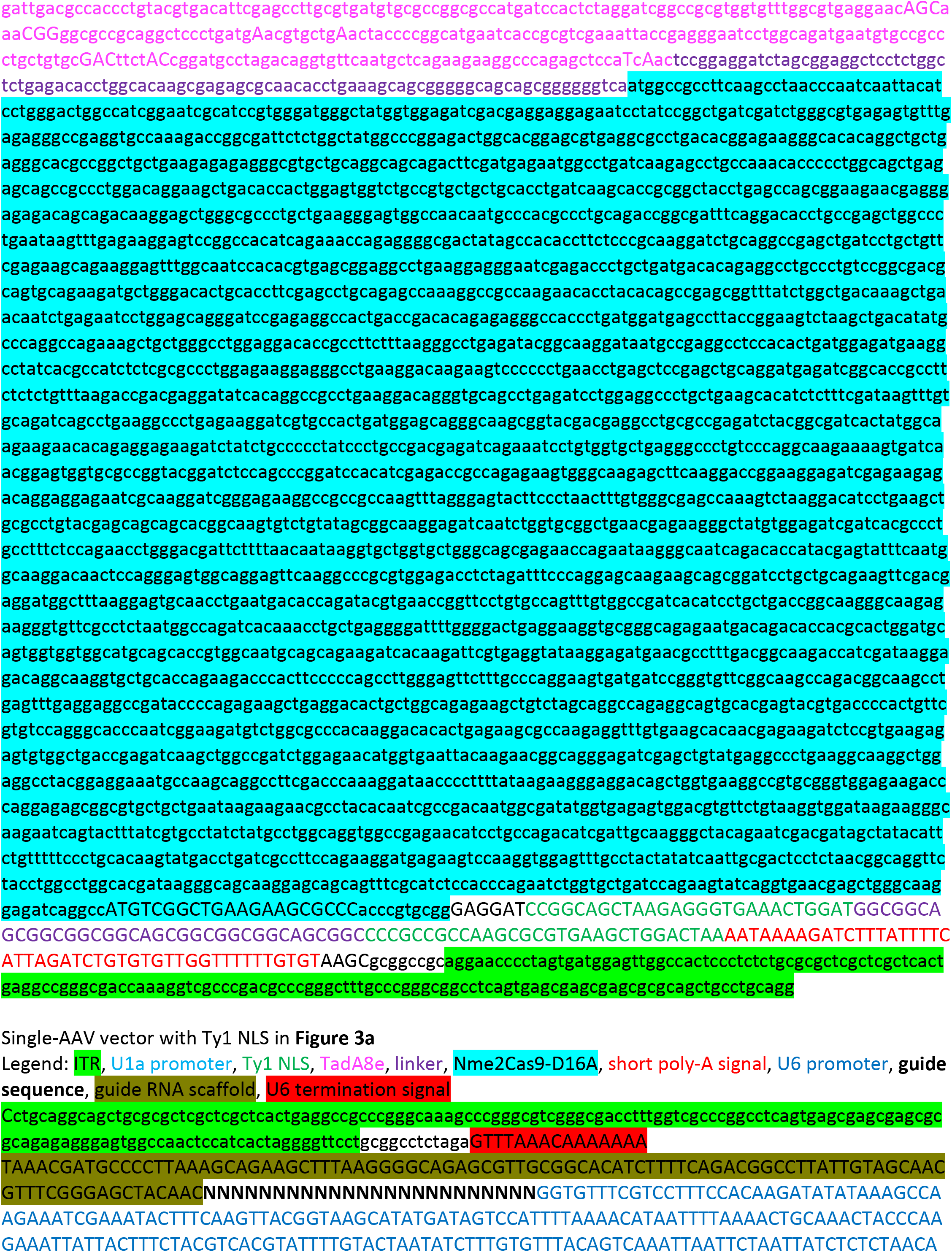

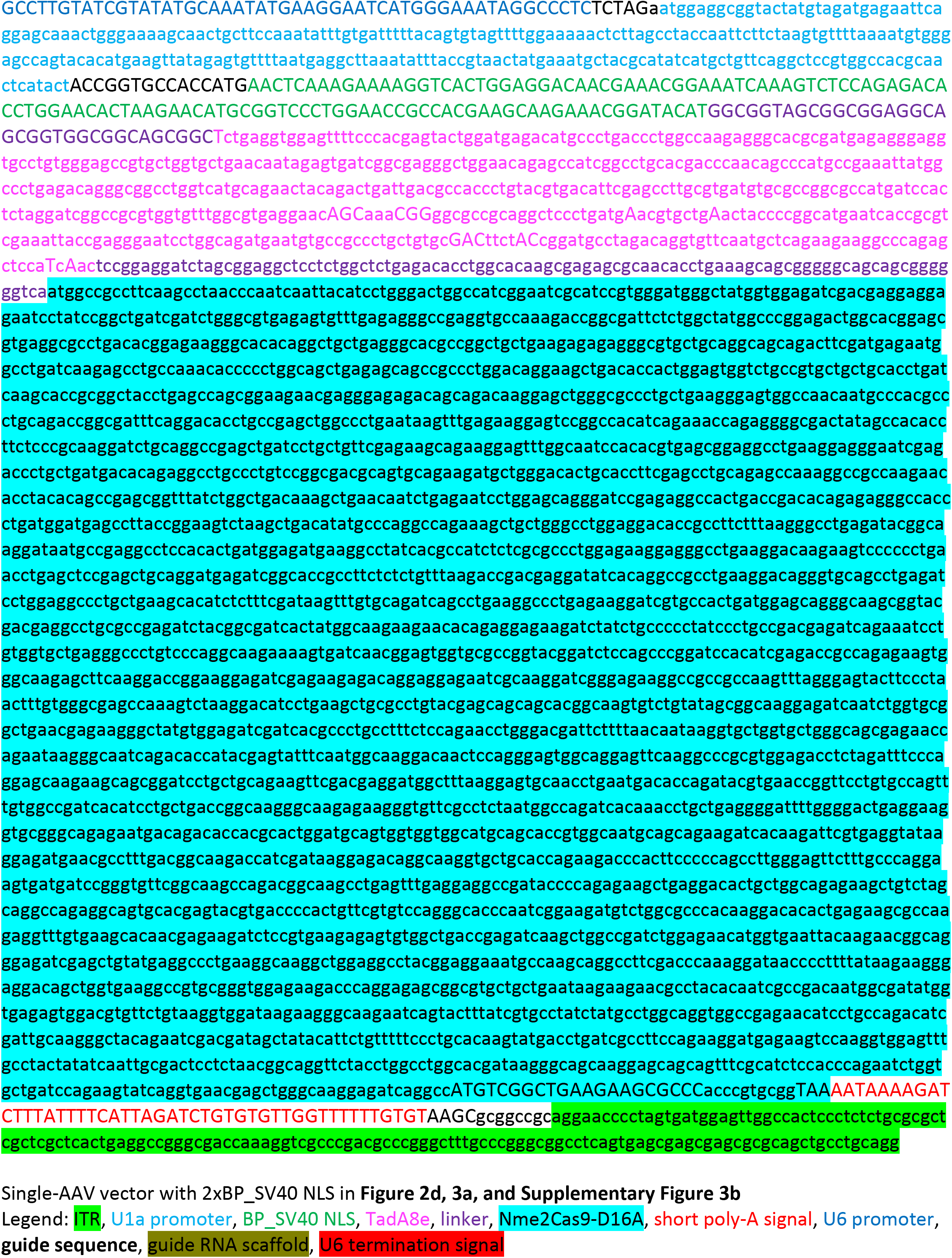

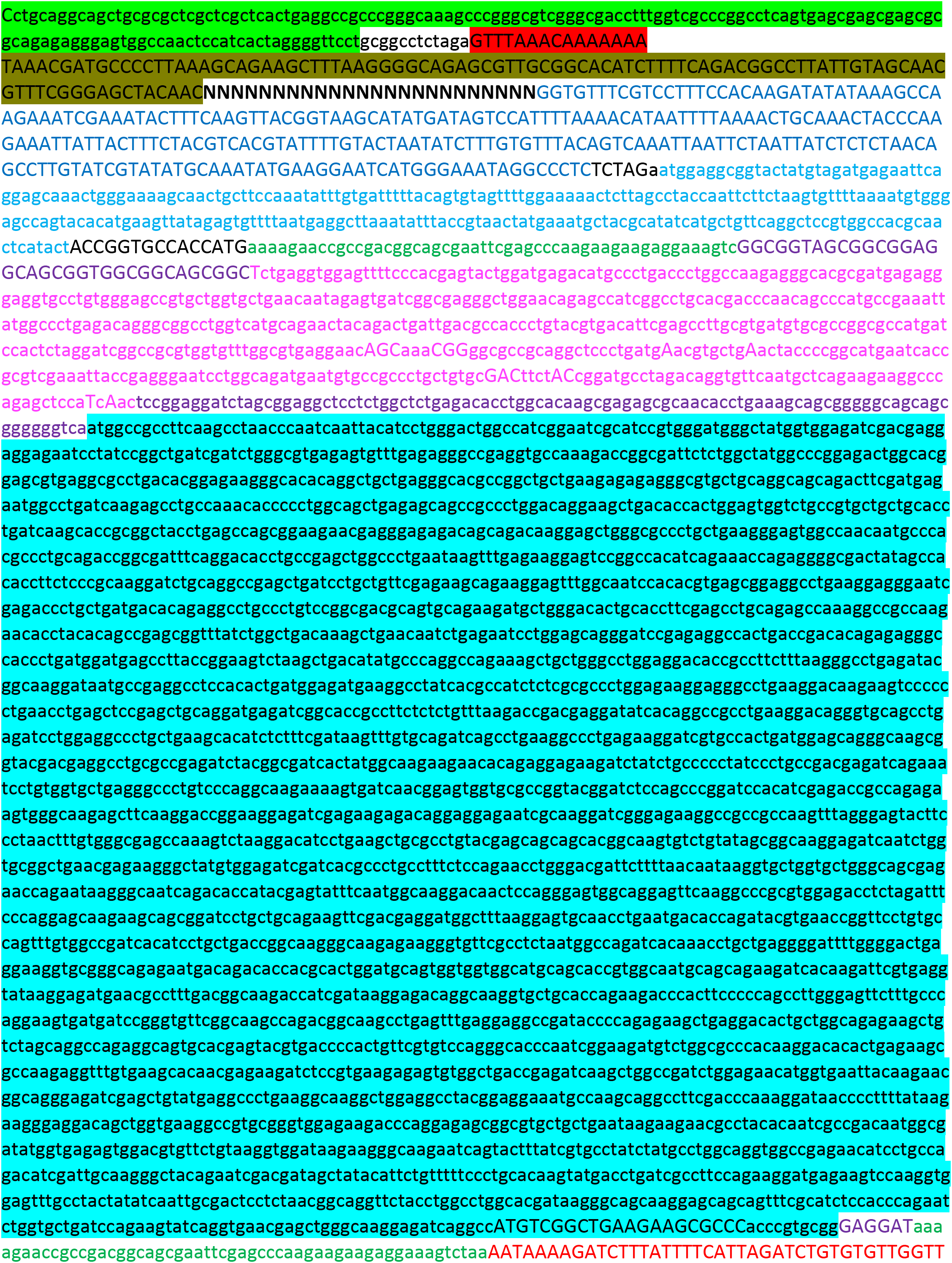

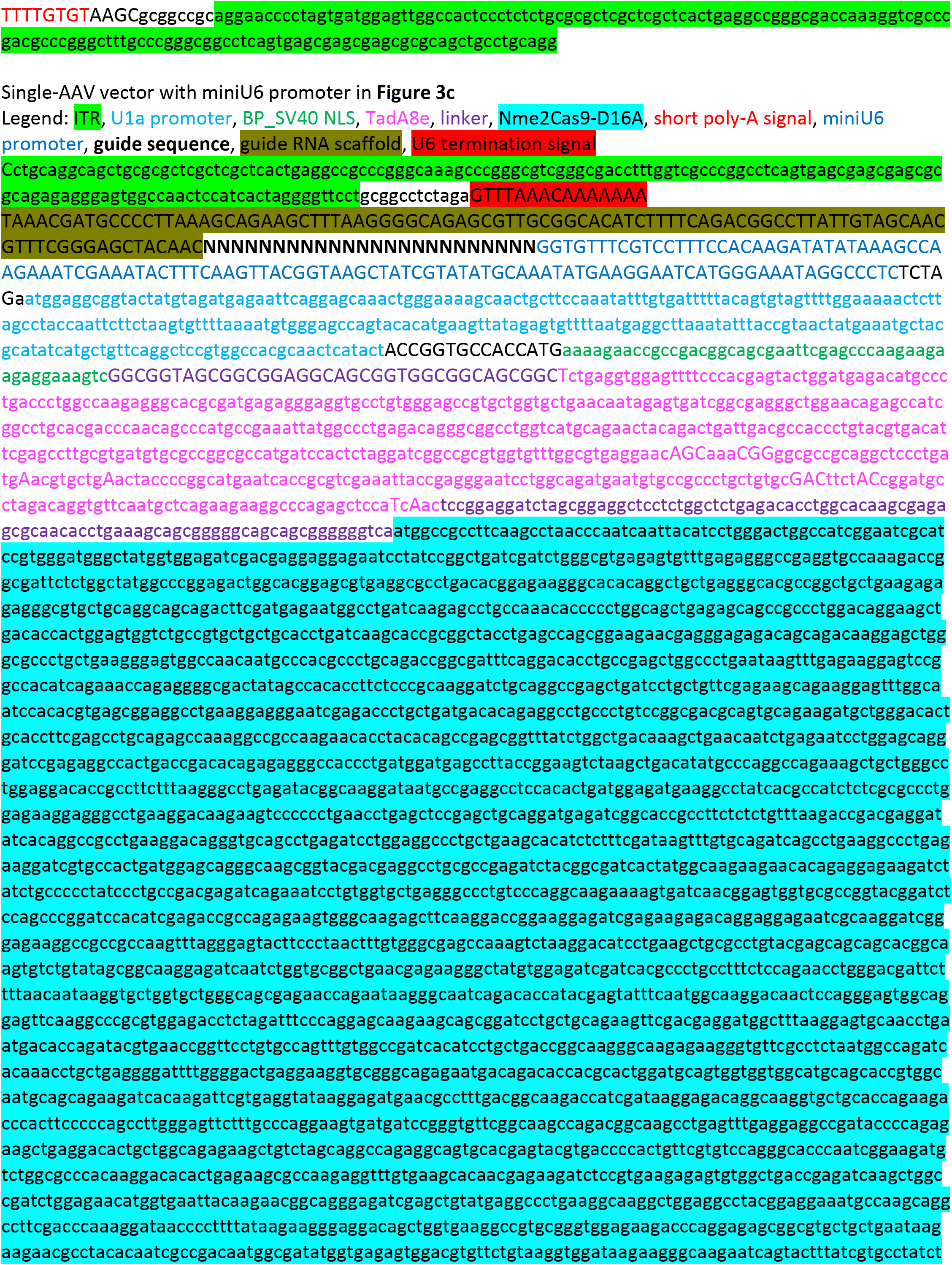

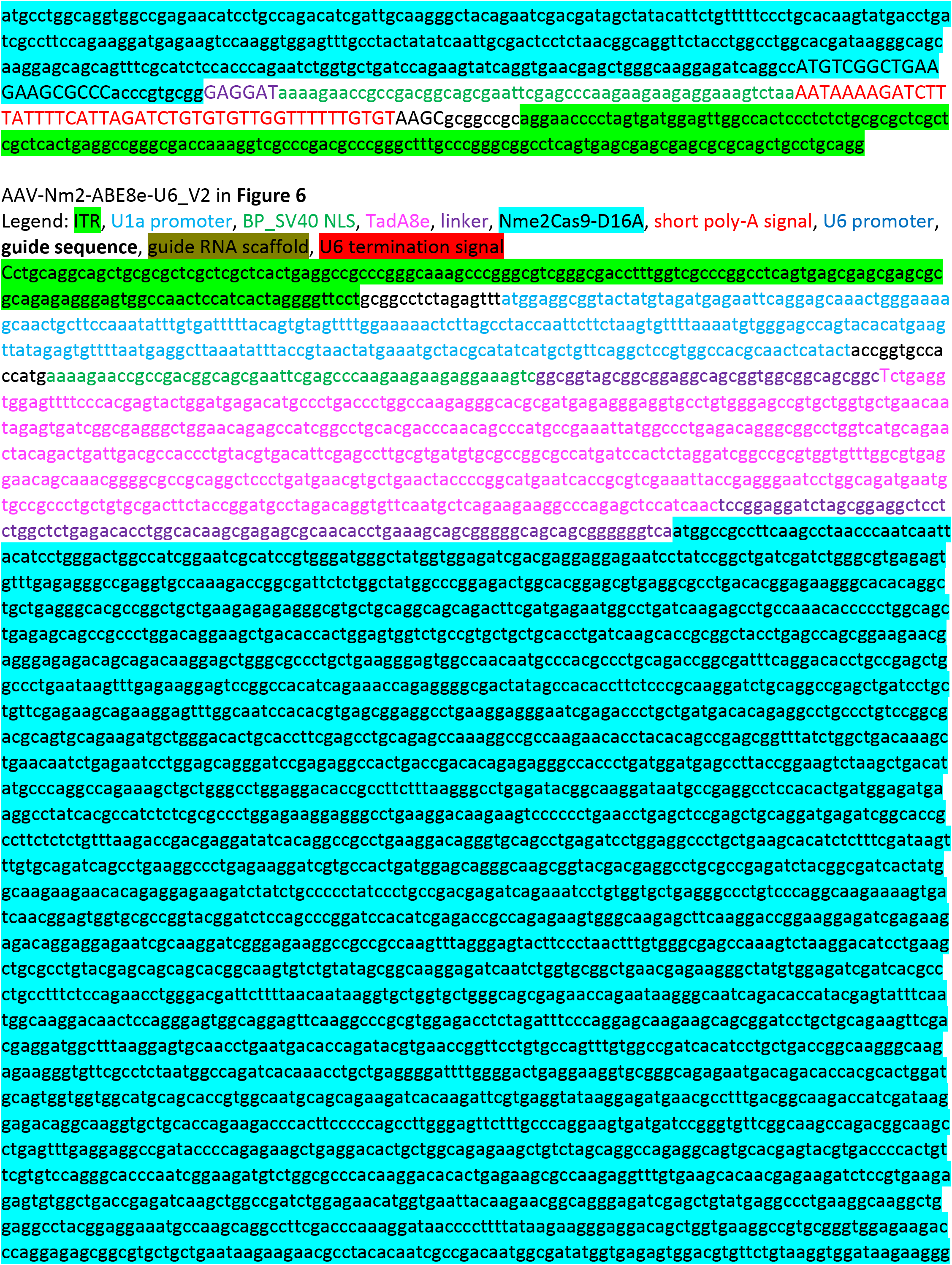

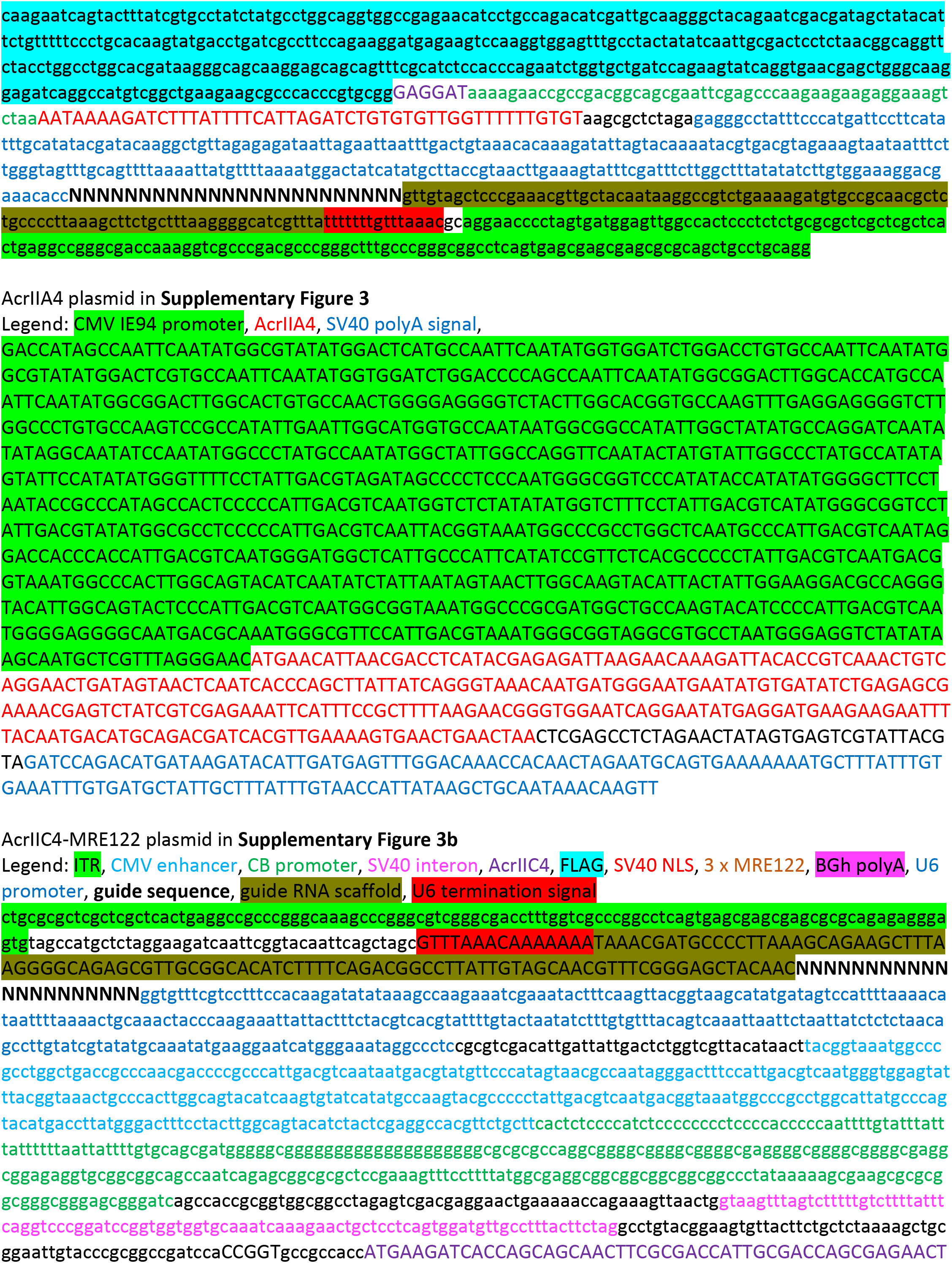

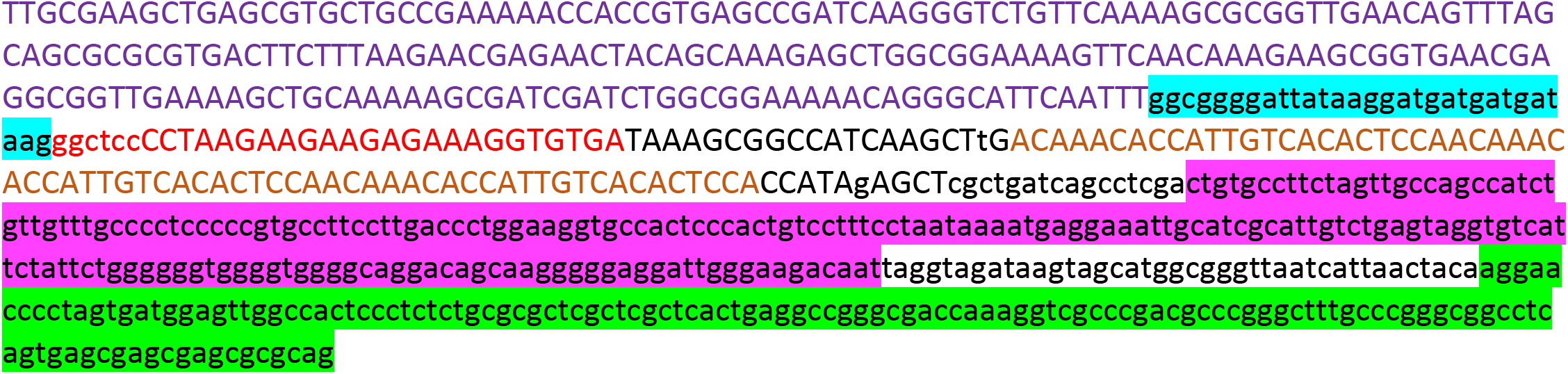

